# Single-cell foundation modeling with species-native protein tokens links regenerative competence across frog and mouse

**DOI:** 10.64898/2026.07.27.740807

**Authors:** Hu Cang, Sha Sun

## Abstract

Appendage regenerative capacity varies dramatically across species, developmental stages, and anatomical sites, yet comparing functional transitions across organisms remains difficult because gene vocabularies diverge. Cross-species single-cell analysis conventionally collapses divergent genomes to one-to-one orthologs—a reduction that is not neutral. Here, we construct a species-native input representation for allotetraploid *Xenopus laevis* that preserves duplicated L and S homeologs (96.39% feature coverage versus 53.14% under symbol collapse) within a frozen universal cell embedding (UCE). A causally validated tail-organizer contrast defines a portable vector competence ruler. While baseline representations (direct expression, SVD, Harmony) recover organizer identity, only species-native UCE preserves the stage-52-versus-stage-58 limb competence transition, which ortholog collapse reverses. Applied without refitting, the ruler distinguishes regenerative from fibrotic digit repair in adult mice and resolves an aligned component in state-balanced macrophages, an ordering reproduced by simpler representations. Preserving species-native gene vocabularies carries functional contrasts across evolutionary and genomic boundaries.

## Introduction

Regenerative competence varies dramatically across tissues, developmental stages, and anatomical sites.^1–4^ Within a single organism, appendage regeneration can vanish during maturation or vary across millimeters of the same anatomical structure. A young *Xenopus laevis* tadpole regenerates a fully patterned limb that an older tadpole cannot, whereas an adult mouse regenerates a distal digit tip but heals proximal amputations by scarring.^5–14^ These functional transitions define competence as a contextual biological state revealed through contrasts between regenerative and refractory conditions. Transporting such an experimentally anchored contrast across species boundaries to identify concordant repair compartments represents a central challenge in comparative biology.

Single-cell transcriptomics provides a powerful foundation for cross-species cell comparison, yet conventional integration frameworks—such as SAMap, SATURN, and Harmony—focus primarily on cell-identity alignment by warping transcriptomic manifolds to co-cluster equivalent cell types.^15–20^ Matching cell classifications, however, does not reveal whether tissues share functional capacity: related signaling centers exist in both competent and refractory tissues, while equivalent repair outcomes are frequently executed by distinct cellular lineages across species. Comparative biology requires a complementary paradigm: freezing a causally validated biological contrast in one organism as a directional vector ruler, and asking where a concordant outcome difference appears across cell compartments in another.

The mathematical foundations of this approach build on established vector-displacement principles, directional comparisons, and frozen reference mapping, as exemplified by scGen and Symphony.^21–24^ Universal cell embedding (UCE) supplies a unified cross-species coordinate space.^25,26^ We combine these principles into a frozen foundation ruler: the unit-normalized difference between population centroids of two experimentally defined biological states. Applied without refitting, the ruler provides a common direction along which prespecified differences between library-level population summaries can be evaluated without target-label fitting. UCE’s protein-derived gene tokens make this anchored contrast portable across divergent and duplicated gene vocabularies without requiring one-to-one ortholog restriction.

Appendage regeneration offers an ideal sequential test for this framework. First, surgical ablation and transplantation experiments in the *X. laevis* tail established regeneration-organizing cells (ROCs) as an essential signaling center, anchoring a ROC-versus-ordinary-epidermis ruler.^27^ We froze that direction and tested it in an independent limb atlas, where it recognized the related apical ectodermal ridge (AER) program and tracked competence loss between stages 52 and 58.^5^ Second, we froze the stage-52-versus-stage-58 limb contrast and transported it into mouse digit repair, where distal P3 regenerates and proximal P2 heals by fibrosis.^6–14^ Preserving allotetraploid *X. laevis* L and S homeologs in UCE enabled both transfers without collapsing the frog data to one-to-one orthologs. The mouse outcome ordering survived source-composition and target-depth controls and recurred within state-balanced macrophage repair populations; external skin and gene analyses then tested developmental breadth and molecular interpretation. Together, these results establish a group-level, cross-species test of functional-contrast transport.

## Results

### Species-native protein tokens place allotetraploid *Xenopus* in frozen foundation space while preserving homeolog identity

To apply foundation embeddings to an unstudied polyploid organism, we first addressed a genomic hurdle: *Xenopus laevis* is an allotetraploid carrying distinct L and S homeologs, and it was absent from UCE pretraining.^25,28^ Rather than collapsing homeologs onto a diploid reference, we constructed a species-native input asset that preserves L/S identity across nearly all assayed features as the species enters the frozen model.

We linked each assayed *X. laevis* gene to its encoded protein and genomic position. Following published ESM2, UCE and SATURN workflows, we averaged the amino-acid representations within each protein to obtain one protein vector.^19,25,26^ This species-native route represented 30,396 of 31,535 assayed features (96.39%; Fig. 1a) while retaining L and S homeologs as separate protein-informed tokens. The UCE model weights remained unchanged, and tail and limb cells were embedded independently in two sampling runs with different random gene draws (seeds 42 and 43).

**Figure 1.**
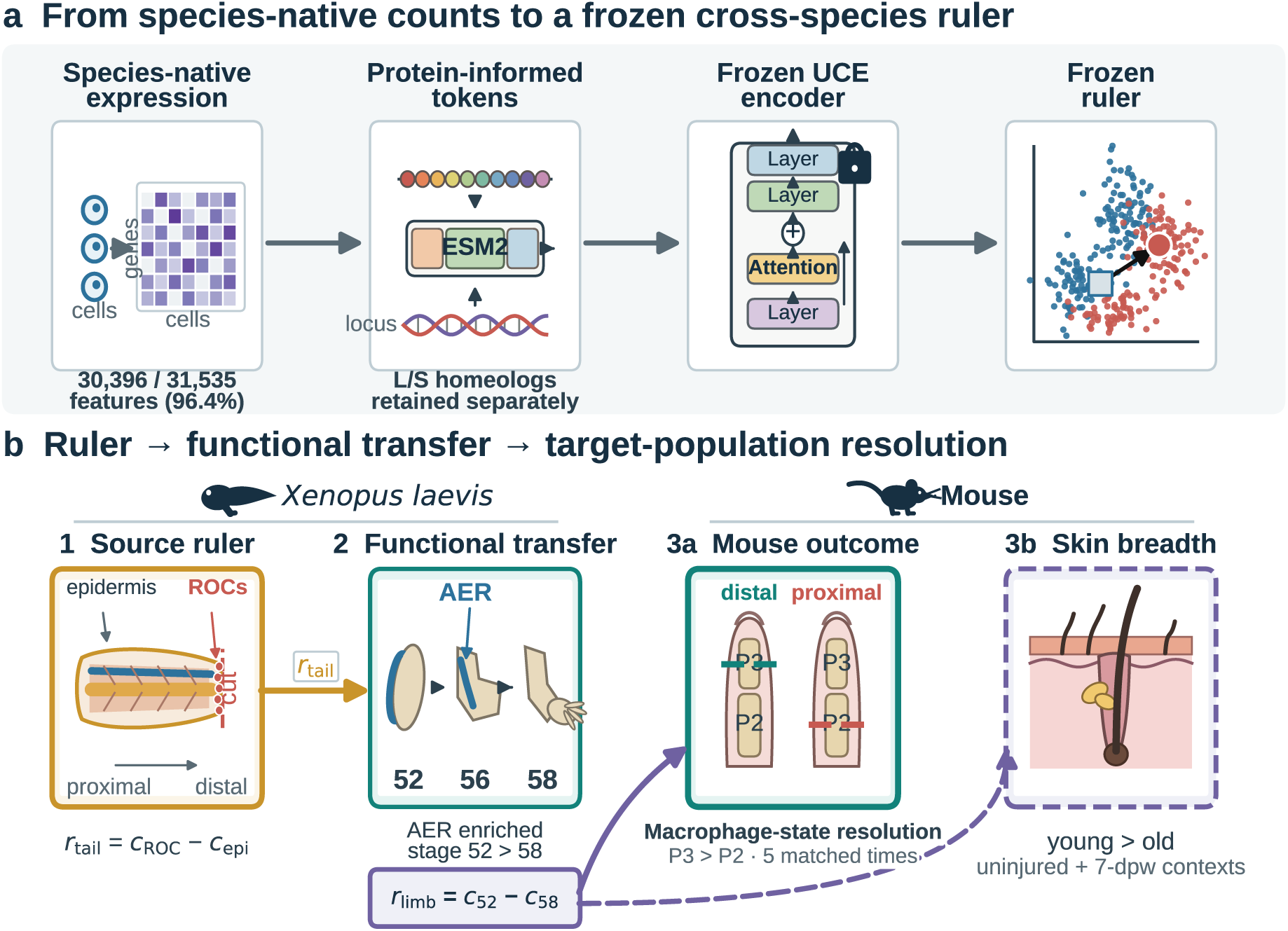
A frog tail organizer contrast becomes a frozen cross-species ruler. a, Species-native *X. laevis* features, ESM2-15B residue-mean protein representations and genomic loci enter frozen UCE. The species-native input covers 30,396 of 31,535 features (96.39%) while retaining L/S homeologs; an equal-library group contrast defines the ruler. **b,** A tail ROC-minus-epidermis ruler tests organizer identity and developmental competence in an independent limb atlas. The limb stage-52-minus-stage-58 ruler then searches mouse digit injury, where the aligned P3-minus-P2 difference recurs within state-balanced macrophage populations after source-composition and target-depth controls, and tests developmental breadth in skin. Each ruler remains frozen through its downstream test, and deposited libraries provide the inferential units.

For comparison, we built **symbol-collapsed UCE** by mapping L/S features to *X. tropicalis* gene names and summing duplicate mappings. This construction retained 16,759 original features (53.14%) as 10,761 unique symbols and merged 5,998 homeolog entries. Species-native UCE retained 96.39% feature coverage and separate homeolog tokens, and a separate expression analysis examined the retained L/S structure. Fig. 1b summarizes how these embeddings enter the transfer sequence.

### Causally validated tail organizers establish a library-stable reference direction for cross-species transport

To establish a portable ruler grounded in causal biology, we anchored direction construction in a mechanistically validated cell contrast. In the *Xenopus laevis* tail, Regeneration-Organizing Cells (ROCs) form a specialized signaling center whose necessity and sufficiency for tissue regrowth were established through surgical ablation and transplantation experiments.^27^ The deposited source dataset (E-MTAB-7716) contains 254 ROCs and 1,800 ordinary epidermal cells across 14 biological libraries.

Within each library, we calculated unit-normalized centroids for ROCs and ordinary epidermis in UCE space, subtracting the epidermal centroid from the ROC centroid to define a directional vector pointing toward organizer identity. Scaling each library contrast to unit length before averaging the 14 directions ensured that every biological sample contributed equally without cell-abundant libraries dominating (Fig. 2a,b). The resulting consensus direction vector is the **frozen tail-organizer ruler**.

**Figure 2.**
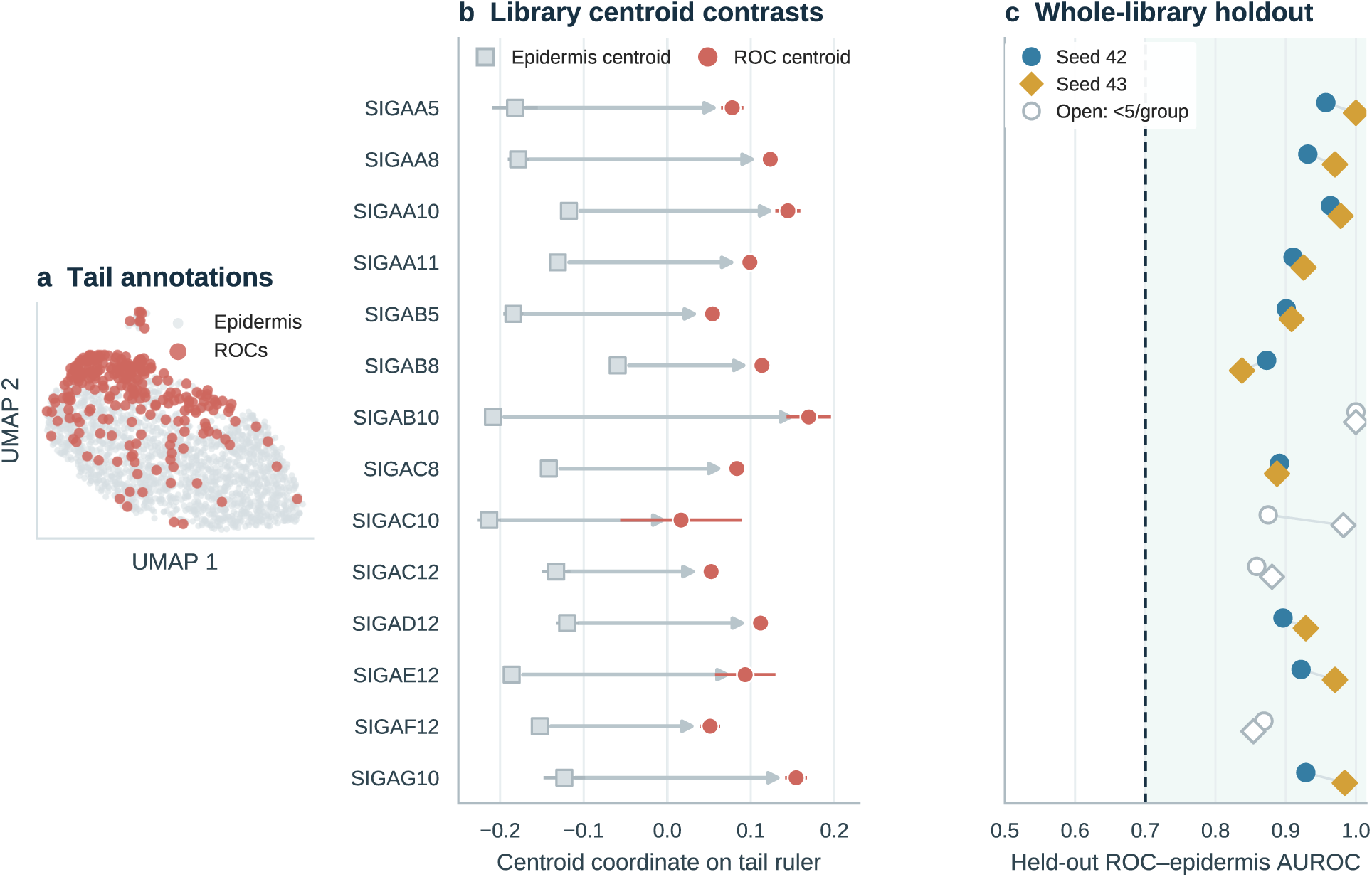
Replicate-level ROC contrasts construct a stable regeneration-organizer ruler. a, UMAP of 2,054 tail epidermal cells coloured by the E-MTAB-7716 ROC annotation. **b,** Unit-normalized ordinary-epidermis (square) and ROC (circle) group centroids from each library, shown at their coordinates along the complete ruler; each connector is the corresponding ROC-minus-epidermis group contrast used in ruler construction. Symbols average seeds 42 and 43; whiskers span them. **c,** Whole-library holdout AUROCs after reconstructing the ruler without each displayed library. Filled symbols meet the prespecified five-cell minimum per group; open symbols provide additional context. The line marks AUROC 0.70. Biological inference uses the library-level group contrasts and whole-library holdouts.

We next asked whether the ruler depended on any one library. For each test, we rebuilt it from 13 libraries and evaluated the omitted library in both UCE sampling runs. We first checked whether omission changed the arrow: a cosine similarity of 1 denotes an identical orientation. We then measured how well the reconstructed ruler separated the two known groups in the omitted library. The area under the receiver operating characteristic curve (AUROC) is 0.5 for chance ordering and 1 for complete separation.

All 14 ROC-to-epidermis arrows pointed in the same direction in both embeddings (Fig. 2b). Every reconstruction remained nearly identical to the complete ruler (minimum cosine 0.9973 and 0.9975 for seeds 42 and 43). It also ranked ROCs above ordinary epidermis in every omitted library with at least five cells per group; even the weakest case reached an AUROC of 0.873 or 0.838 (Fig. 2c; Supplementary Table 2). In that weakest library, a randomly chosen ROC still lay farther toward the organizer side than a randomly chosen epidermal cell in 87.3% or 83.8% of pairings, respectively. The two complete rulers were likewise nearly parallel (cosine 0.9922). The causal ROC contrast therefore produced a stable organizer ruler across libraries and sampling runs.

### A frozen organizer ruler generalizes across tissues to track the developmental loss of limb competence

We next carried the frozen tail-organizer ruler into an independent frog limb atlas. E-MTAB-9104 profiles *X. laevis* hindlimb epidermis across development, including the apical ectodermal ridge (AER), a distal epidermal signaling center that supports limb outgrowth.^5^ We embedded 9,142 Tp63-positive epidermal cells from this atlas independently and applied the ruler unchanged. Within each library containing both AER and other epidermal cells, we asked how far the two groups separated along the tail-defined direction. The AER lay farther toward the organizer side in 12 of 14 libraries in both runs (exact *P* = 5.49 × 10^−4^ and 2.44 × 10^−4^; gaps of 1.25 and 1.58 standard deviations; Fig. 3a,b; Supplementary Table 2). A contrast learned from tail epidermis therefore recognized the related signaling center in independently collected limb tissue.

**Figure 3.**
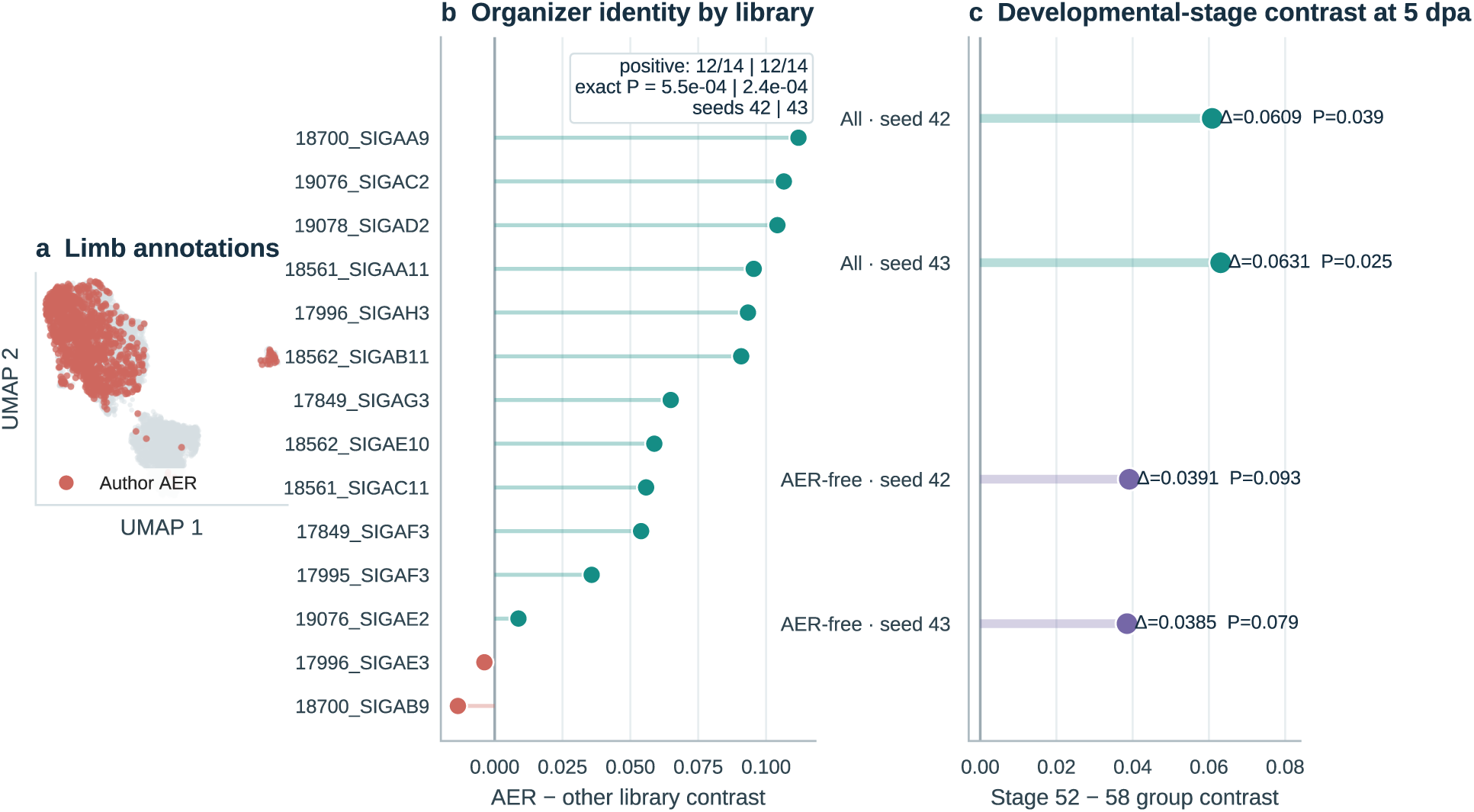
Independent limb data test identity and functional-contrast transfer. a, UMAP of cells annotated as AER in E-MTAB-9104 among 9,142 independently embedded Tp63-positive epidermal cells. **b,** Within each library, the plotted AER contrast is the AER median internal cosine coordinate minus the other-epidermis median; the inset gives positive counts and exact sign-flip probabilities. **c,** The developmental contrast is the equal-library mean of stage-52 library medians minus the corresponding stage-58 mean, shown for all epidermis and after AER exclusion. These raw coordinate differences are unitless and are not probabilities. Stage 56 provides developmental context. Biological inference uses the library-level AER and developmental-stage contrasts.

More importantly, the ruler followed the known developmental loss of limb-regenerative competence. Stage-52 tadpoles regenerate a patterned limb after amputation, whereas stage-58 tadpoles largely form a non-regenerative spike. At five days post amputation, we summarized each library along the ruler and compared the two stages with equal library weight. Expressed relative to between-library variation, stage-52 limbs exceeded stage-58 limbs by 4.23 and 6.52 standard deviations in the two UCE sampling runs (exact *P* = 0.0393 and 0.0250; Fig. 3c; Supplementary Table 2). In raw ruler units, the stage gaps were 0.06085 and 0.06308 (stage-52 means: 0.06218 and 0.06633; stage-58 means: 0.00133 and 0.00325). The transported organizer ruler therefore recognized signaling-center identity and tracked the developmental loss of regeneration.

The early-stage signal also extended beyond annotated AER cells. After removing those cells, approximately 61–64% of the full stage contrast remained (stage-52-minus-stage-58 gaps of 0.0391 and 0.0385 ruler units; *P* = 0.0929 and 0.0786; Fig. 3c), consistent with a broader epidermal component beyond the annotated AER population. Repeating the embeddings with ten different random gene samplings reinforced the transfer: all 100 tail-by-limb run combinations retained a positive stage-52-minus-stage-58 difference, and the average gap was 0.0611 (*P* = 0.0339; Supplementary Fig. 4). Together, the AER and stage results establish the tail-to-limb step: the frozen organizer ruler recognized a related signaling center and retained the developmental competence contrast.

### The vector competence ruler decouples functional state transport from cell-identity matching

Recognizing an organizer and preserving its regenerative state are different tasks. We therefore evaluated each representation twice: AER recovery tested cross-tissue identity, whereas stage-52-versus-stage-58 separation tested whether the functional contrast survived transfer. Alongside species-native and symbol-collapsed UCE, we compared three familiar alternatives built from 9,960 shared gene symbols: direct expression, a lower-dimensional SVD representation learned from tail data, and Harmony integration.^16,29^ Every method was evaluated on the same biological groups, with each experimental library given equal weight.

On the competence test, species-native UCE produced stage gaps of 4.23 and 6.52 on a scale where 1 equals one pooled between-library standard deviation (exact *P* = 0.0393 and 0.0250; Fig. 4a). Direct expression, tail-trained SVD and Harmony produced much smaller gaps of 0.22, 0.41 and 0.23 on the same scale (*P* = 0.438, 0.382 and 0.423), while symbol-collapsed UCE reversed the contrast in both runs (−3.73 and −0.70). Species-native UCE thus preserved the regenerative transition that the other representations did not.

**Figure 4.**
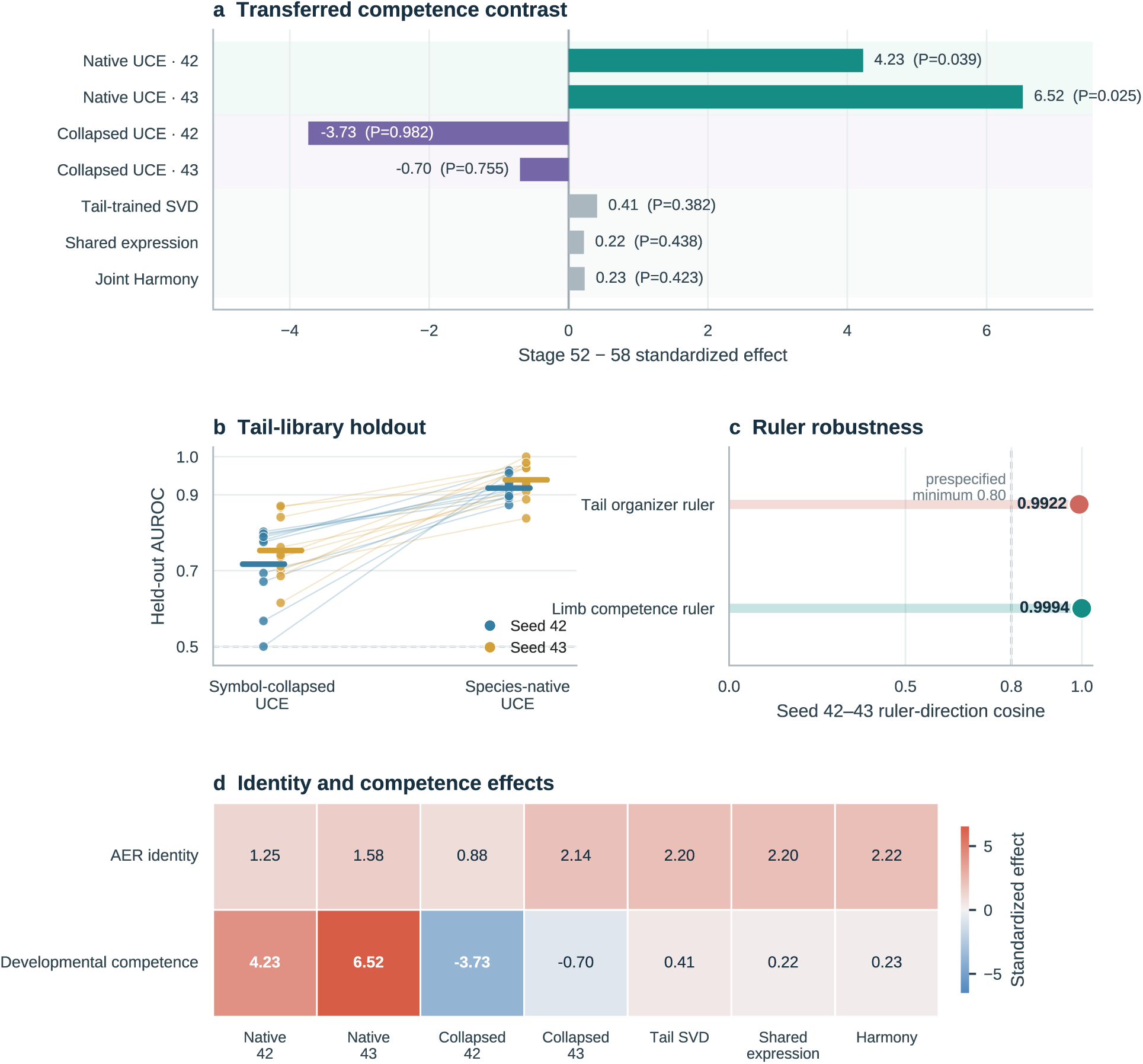
Species-native UCE preserves competence beyond organizer identity matching. **a,** Standardized stage-52-minus-stage-58 effects—raw group differences divided by pooled between-library standard deviations—and exact stage-label probabilities. Direct expression and tail-trained SVD used 9,960 symbols shared with GSE165816 and were applied unchanged to limb; Harmony jointly fitted tail, limb and unlabelled human epithelial expression. Human outcomes remained held out. **b,** Paired symbol-collapsed and species-native UCE AUROCs after leaving out each eligible tail library; horizontal strokes mark equal-library means. Species-native UCE exceeded symbol-collapsed UCE in 10/10 libraries per seed (*P* = 9.77 × 10^−4^). **c,** Seed-42-versus-seed-43 cosine similarities for organizer and competence rulers; the line marks 0.80. **d,** Standardized AER-identity and developmental-competence effects on the same scale as panel a. All representations recover organizer identity; only species-native UCE additionally preserves competence. This panel carries the representation-dependence claim of the study; the mouse endpoint of Fig. 5 does not (Supplementary Fig. 10). Biological inference uses group contrasts at the deposited-library level.

A separate tail-library check asked whether preserving the native frog gene vocabulary strengthened recovery of the causal ROC contrast. We left out one complete library at a time, rebuilt the ruler and evaluated it in the omitted library. Species-native UCE exceeded symbol-collapsed UCE in all ten libraries that met the five-cell-per-group minimum in each run, increasing mean AUROC from 0.717–0.753 to 0.917–0.939 (paired-sign *P* = 9.77 × 10^−4^; Fig. 4b). It also retained 96.39% of assayed features and separate L/S homeolog tokens, compared with 53.14% feature coverage after symbol collapse. Thus, retaining the native frog feature space both preserved more measured biology and strengthened recovery of the causal organizer contrast (Supplementary Fig. 3). The tail-organizer and limb-competence directions were nearly identical across sampling runs (cosines 0.9922 and 0.9994; Fig. 4c), and the limb library-summary contrasts were likewise strongly concordant (Spearman ρ = 0.881).

All methods nevertheless recognized AER identity. Every limb library containing both populations showed a positive AER-versus-other-epidermis contrast with direct expression, tail-trained SVD and Harmony (standardized effects 2.20, 2.20 and 2.22; exact *P* = 6.10 × 10^−5^ each; Fig. 4d), while species-native UCE also produced a positive AER effect (standardized effects 1.25 and 1.58 across 12 of 14 libraries). Yet only the species-native UCE route preserved the developmental competence transition in this comparison. This result separates identity recovery from competence preservation within frog and establishes the representation used for the subsequent frog-to-mouse test.

### The frozen frog competence ruler distinguishes regenerative and fibrotic outcome groups in adult mouse digit repair

Having established a frog competence ruler, we asked where a related outcome contrast appears in adult mouse digit repair. Distal P3 regenerates, whereas proximal P2 heals by fibrosis.^6–14^ We applied the ruler unchanged and compared P3 and P2 libraries collected at matched times after amputation. In the six-library primary series (GSE135985), every regenerative P3 library lay above every fibrotic P2 library, and the P3 group exceeded P2 at both collection times (Fig. 5a,b). This complete 3-versus-3 separation gave the largest statistic among the nine valid within-time label allocations in both UCE runs. The design permits only those nine allocations, so rank 1 corresponds to the finest attainable one-sided exact probability, 1/9 = 0.111. The time-adjusted P3-minus-P2 gaps were 0.0952 and 0.0956 in raw ruler units. Cell-level coordinates were strongly concordant between UCE sampling runs in both dataset series (Supplementary Fig. 1). The same initial tissue-wide ordering appeared at days 12, 17 and 28 in a second dataset series, extending the search across later repair times.

**Figure 5.**
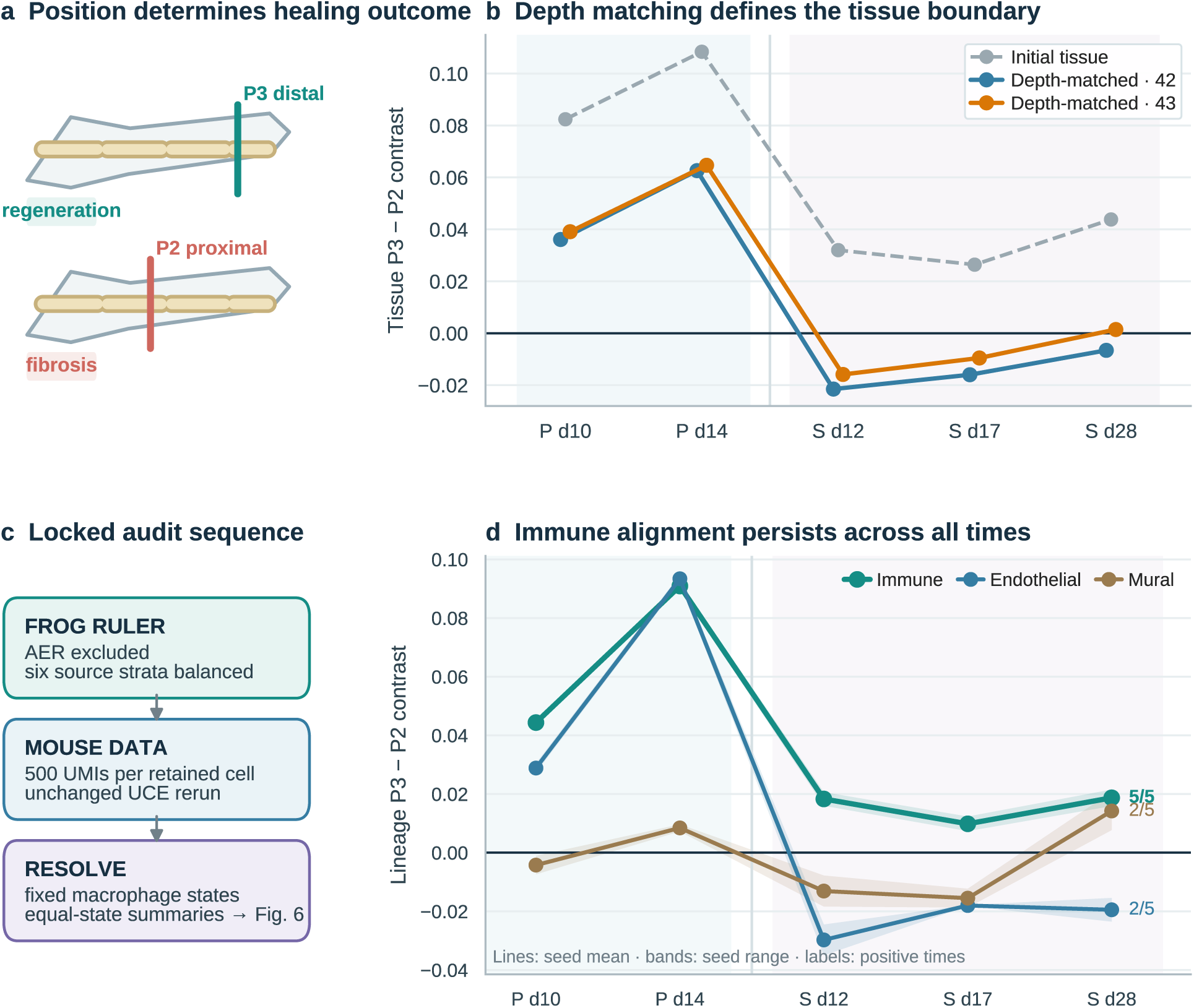
The frozen frog ruler searches mouse tissue and identifies the aligned lineage. **a,** Distal P3 regenerates; proximal P2 scars. **b,** Initial (grey) and AER-free, depth-matched (coloured) P3-minus-P2 contrasts. Each library is summarized by its median internal cosine coordinate, and each point is the mean P3 summary minus the mean P2 summary at one matched time. P denotes the matched primary series and S the cross-study second series. Depth matching preserves both primary contrasts and delineates the tissue-wide scope across later times. **c,** Analysis sequence from frog composition balancing and mouse depth matching to fixed macrophage-state summaries. **d,** Controlled screen of lineages represented by at least 20 cells from each outcome at all five matched times. Lines show seed means, bands span seeds and right labels give the number of positive times. Immune cells are positive at all five times in both UCE runs. Biological inference uses P3-minus-P2 contrasts between deposited-library summaries. The six-library design admits nine within-day label allocations, so its finest attainable exact probability is 1/9 = 0.111; direct expression, source-only SVD and Harmony reach the same endpoint, so this ordering is reported as representation-general rather than as evidence of UCE superiority (Supplementary Fig. 10).

Because this design permits only nine within-time label allocations, its finest attainable one-sided exact probability is 1/9 = 0.111, and the endpoint is correspondingly coarse for discriminating between representations. We tested that directly by running the complete frog-to-mouse chain separately for direct expression, source-only SVD and Harmony, each constructing its own frozen ruler from the same AER-free, composition-balanced frog libraries and applied to the same depth-rarefied mouse cells. All three reached the same endpoint as species-native UCE—rank 1 of 9, complete 3-versus-3 separation and five of five source leave-one-out positives—and source-only SVD produced the largest raw contrasts of any representation tested (Supplementary Fig. 10, Supplementary Table 4). The mouse ordering is therefore representation-general. We report it as evidence that the transported biological difference is robust to the choice of representation, not as evidence that UCE is uniquely required at this endpoint. The representation-dependent step is the frog competence contrast of Fig. 4, where the same three comparators fail.

### Pre-inference depth rarefaction and source balancing confirm that cross-species alignment survives technical controls

We next placed the mouse result under a deliberately stricter analysis that equalized the two main technical imbalances: frog source-cell composition and mouse sequencing depth (Fig. 5c). We removed AER cells from the frog source, balanced six epidermal and cell-cycle groups, and reduced every retained mouse cell to exactly 500 raw unique molecular identifier (UMI) counts before rerunning the embedding and transfer from the beginning.

P3 remained above P2 at both matched primary times, with the observed label assignment again ranking first of nine (exact *P* = 1/9). Approximately half (52–54%) of the original tissue-wide separation remained under these strict controls (equal-time mean gaps of 0.0494 and 0.0519 ruler units; day 10: 0.0361 and 0.0391; day 14: 0.0626 and 0.0647; Fig. 5b; Supplementary Fig. 6). When the three later cross-study comparisons were added, the total was two of five and three of five positive time points in the two runs (Fig. 5b; Supplementary Fig. 6). This narrower tissue-wide pattern focused the next analysis on the cell compartment in which the alignment persisted.

### Cross-species competence alignment resolves a repeatable component in macrophage repair states

The changing whole-tissue result suggested that the aligned biology might reside in one cell compartment rather than across the entire digit. We therefore applied the same depth-matched analysis to broad lineages defined without outcome labels. Lineage selection was then outcome-informed: among lineages represented by at least 20 cells in every P2 and P3 library at all five matched times, immune cells were the only lineage with P3 above P2 at all five times in both UCE runs. Endothelial and mural lineages were positive at two of five times in both runs (Fig. 5d). This controlled screen nominated immune cells for finer analysis.

Within the immune compartment, predefined marker programs identified macrophage-like cells and divided them into lipid/phagocytic, resident/remodelling, inflammatory and interferon states. To keep state composition comparable, a state entered the analysis only when every library in that series contained at least ten assigned cells. Two states qualified in the primary series and three in the second (Supplementary Fig. 7a), and each qualified state contributed equally to its library summary.

Across both dataset series and all five matched times, P3 exceeded P2 in both sampling runs (five-time means of 0.01765 and 0.02217 ruler units; contrasts ranging from 0.000150 to 0.0465 in the first run and 0.00535 to 0.0519 in the second; Fig. 6a). Because each macrophage state contributed equally within a library, the ordering reflects shifts within matched state categories rather than differences in their abundance. The aligned P3-minus-P2 difference was therefore reproducibly resolved within state-balanced macrophage repair populations.

**Figure 6.**
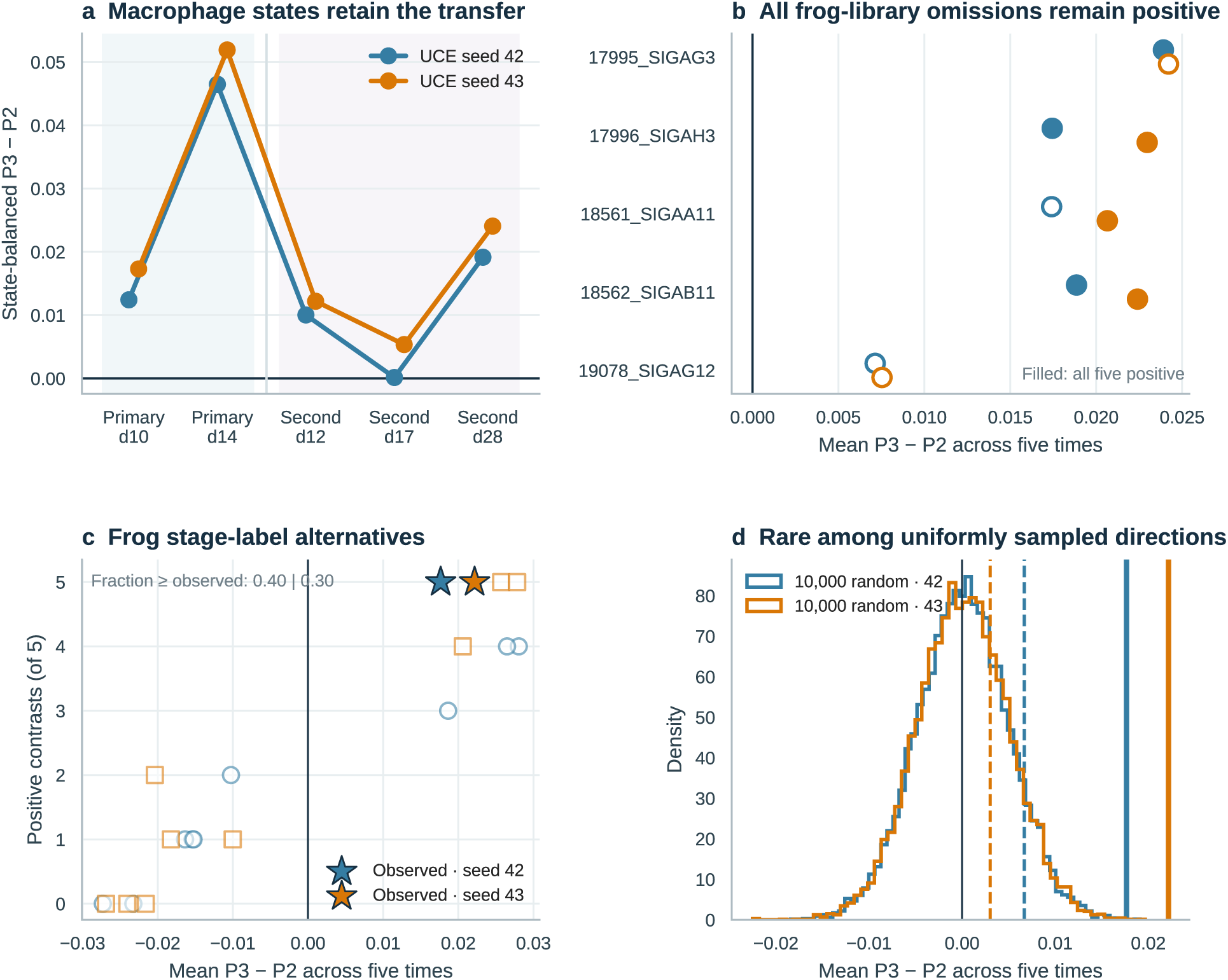
The frozen competence ruler resolves a repeatable aligned component in macrophage repair states. **a,** Each library summary is the equal mean of eligible state-specific median internal cosine coordinates; primary and second series use two and three eligible states. Each plotted value is the mean P3 summary minus the mean P2 summary at one matched time, and all five are positive in both seeds. **b,** Unweighted means of the five time-specific raw contrasts from five leave-one-frog-library-out rulers per seed; all remain positive. **c,** The same five-contrast mean and positive-sign count for all ten allocations of three early and two late labels across the five frog libraries; stars mark the biological allocation. **d,** The observed five-contrast mean against 10,000 uniformly sampled directions; dashed lines mark cell-cycle controls. A mean raw contrast this large is rare (*P* = 2.0 × 10^−4^ and 1.0 × 10^−4^), whereas approximately 12.5% of random directions yield five positive signs.

### Source-label and random-direction controls define ruler specificity

We first asked whether the result depended on any single frog library. Omitting each of the five stage-52 or stage-58 libraries in both sampling runs produced ten reconstructed rulers; all retained a positive five-time mouse mean and remained aligned with the complete ruler (cosine 0.794–0.996; Fig. 6b). We next rebuilt the ruler for every possible division of the five frog libraries into groups of three and two. The biological stage allocation produced P3 above P2 at all five mouse times in both runs and was the only allocation to satisfy that criterion in both. Considered separately, one of ten and three of ten allocations yielded five positive contrasts, while four of ten and three of ten produced a mean at least as large as the biological allocation (Fig. 6c). The five-library source design therefore supports cross-run directional consistency, while its resolution for distinguishing the observed stage labels from every alternative allocation remains limited.

We then compared the ruler with 10,000 directions sampled uniformly across the full UCE space. Only one and zero sampled directions produced a state-balanced macrophage mean as large as the observed ruler (+1-corrected empirical *P* = 2.0 × 10^−4^ and 1.0 × 10^−4^; Fig. 6d). A biological control built from cycling-versus-G1 frog epidermis was also smaller (means 0.00668 and 0.00301), 2.6-fold and 7.4-fold below the competence ruler. Because the immune compartment had been selected from the lineage screen, we repeated that selection rule for every random direction. None of 10,000 uniformly sampled directions selected a complete lineage with five positive contrasts and a mean as large as the observed immune result in either run (+1-corrected *P* = 1.0 × 10^−4^; Supplementary Fig. 9b).

Uniform directions do not capture the anisotropic geometry actually occupied by frog source populations. We therefore added a post hoc sensitivity in which random directions were restricted to the source-supported covariance structure estimated from 30 AER-free frog stratum-by-library centroids. Five source principal components retained 91.7% and 93.2% of this variation (Supplementary Fig. 9a). After repeating lineage selection for every sampled direction, the observed immune result lay in the upper tail of this more demanding null (*P* = 0.0820 and 0.0382); with the macrophage summary fixed, the corresponding probabilities were 0.0970 and 0.0478 (Supplementary Fig. 9b,c). These source-geometry results complement, rather than replace, the exhaustive source-label control and show how the evidence changes as the null moves from arbitrary directions to directions supported by frog source variation.

As a post hoc reciprocity check, we applied a mouse-derived ruler back to frog limb libraries. We built this ruler from the depth-matched primary series, giving lipid/phagocytic and inflammatory macrophage states equal weight within each library and isolating the P3-versus-P2 direction after accounting for collection day. Applied unchanged to frog, the mouse-derived ruler achieved a strict stage-52 > stage-56 > stage-58 stage-mean ordering across the eight limb libraries in both runs (exact *P* = 0.00536 and 0.00357; stage-52-minus-stage-58 gaps of 0.0313 and 0.0281 ruler units; Supplementary Fig. 8a,b). After AER removal and six-stratum epidermal balancing, the gaps remained 0.0283 and 0.0302 (Supplementary Fig. 8c). The same ruler placed tail ROCs above ordinary epidermis in 11 of 14 and 13 of 14 libraries (exact *P* = 0.00330 and 1.83 × 10^−4^; Supplementary Fig. 8d). All six leave-one-mouse-library-out rulers retained a positive frog stage gap, and the observed mouse outcome assignment ranked first and second among the nine within-day allocations (Supplementary Fig. 8e). This reciprocal construction provides a post hoc consistency check on the geometry of the forward alignment.

### External skin data extend the developmental contrast across uninjured and wounded tissue

To test whether the competence ruler captures a broader developmental contrast in skin and hair follicles, a prespecified external-dataset analysis examined GSE153596.^30^ This study includes unwounded skin collected at postnatal day 2 (PND2) or PND21, together with wounds initiated at PND2 or PND21 and collected seven days later. Among the wounded samples, all three libraries wounded at PND2 exceeded all three libraries wounded at PND21 in both UCE runs, ranking first among 20 possible age-label assignments (exact *P* = 1/20 = 0.05; contrasts of 0.0382 and 0.0411 ruler units; Supplementary Fig. 5a,b).

The young-above-old ordering also appeared in unwounded skin (contrasts of 0.0461 and 0.0474 ruler units; Supplementary Fig. 5c–e). Because the wounded groups were collected at approximately PND9 and PND28, whereas the unwounded groups were collected at PND2 and PND21, their difference of contrasts is descriptive rather than an isolated injury effect. The concordant ordering across both contexts nevertheless extends the ruler-defined developmental contrast to cutaneous tissue.

### A focal lipid-handling and matrix-remodelling module interprets the transported contrast

To interpret the transported contrast, we compared frog epidermal and mouse immune gene effects across 11,549 shared symbols. Genome-wide rank concordance was low (ρ = −0.090 to 0.128; Supplementary Fig. 2), indicating that signaling epidermis and macrophages implement the aligned outcome in distinct transcriptional contexts. Within this broad divergence, the shared molecular signal concentrated in a focal tissue-remodelling program.

Against this divergent background, a focused post hoc test of six repair-, lipid- and phagocyte-related genes found four shared-positive genes where 0.82 were expected (4.85-fold enrichment; *P* = 0.00423; Bonferroni-adjusted *P* = 0.00846; Supplementary Table 3). Because this repair set overlaps the predefined macrophage programs, the enrichment provides molecular interpretation of the resolved compartment. The broader focal bridge included *Apoe*, *Lpl*, *Lum*, *Psap*, *Cst3* and *Ctsb* (Supplementary Fig. 2), linking the transported contrast to tissue remodelling across different cell types.

## Discussion

Biological contrasts defined in one organism can serve as portable vector rulers for locating concordant functional states across species boundaries. In this study, we established a progressive cross-species transfer framework. First, causally validated amphibian tail organizers anchored a baseline reference vector. Next, an independent limb atlas demonstrated that this vector generalizes across tissues to track developmental competence loss. Finally, the resulting frog competence ruler distinguished regenerative P3 from fibrotic P2 mouse digit repair libraries without refitting, resolving a repeatable aligned component within state-balanced macrophage repair populations. Across each stage, foundation embeddings transform complex single-cell transcriptomes into intuitive, directional projections along an experimentally anchored biological axis, enabling zero-shot functional transport across species and gene vocabularies.

This approach bridges foundational concepts in high-dimensional biology: protein-informed latent spaces from ESM2, UCE, and SATURN; vector-displacement modeling from scGen; directional comparison; and frozen reference mapping from Symphony.^19,21–26^ Integration frameworks such as SAMap and SATURN excel at aligning equivalent cell identities across species genomes,^19,20^ but were not benchmarked directly here. Among the representations tested head to head within frog, direct expression, tail-trained SVD, and Harmony all recovered AER cell identity. However, only species-native UCE additionally preserved the developmental competence contrast. That representation dependence is specific to the frog competence step; at the six-library mouse endpoint the same three comparators reproduced the outcome ordering (Supplementary Fig. 10). UCE provides a unified coordinate space in which species-native protein tokens enter independently. This allows the same anchored contrast to cross an allotetraploid frog genome and mammalian gene vocabulary without prior one-to-one ortholog collapse.

Preserving the species-native feature space proved essential for functional transport. The native asset represented 96.39% of assayed *X. laevis* features, retained duplicated L and S homeologs, and substantially improved holdout recovery of the causal ROC contrast relative to symbol-collapsed UCE. The comparison evaluates the complete pipeline—feature representation, embedding geometry, and frozen-ruler construction—rather than isolating a single architectural component. Within this integrated benchmark, species-native UCE preserved both organizer cell identity and the stage-dependent functional contrast that enabled zero-shot transport to mouse.

Extensive control hierarchies clarify the robustness and limits of this cross-species alignment. The competence vector remained remarkably stable across leave-one-out source library omissions. It also survived strict controls that equalized source-cell composition and target sequencing depth. Its macrophage-aligned magnitude proved exceptionally rare among uniformly sampled directions, even when repeating the broad-lineage selection step for every draw. A more demanding, post hoc null restricted directions to the source-supported covariance structure of frog source populations. Under this null, the observed result remained in the upper tail, with selection-adjusted probabilities of 0.0820 and 0.0382 across the two UCE runs. The exhaustive source-label control was less selective by magnitude (4/10 and 3/10 alternatives met or exceeded the observed mean), reflecting the limited statistical resolution of five source libraries. Together, these controls distinguish three key questions: stability to source omission, rarity in embedding geometry, and specificity of the biological stage labels.

These rigorous controls sharpen our understanding of cross-species functional alignment. In adult mouse digit repair, species-native UCE, direct expression, source-only SVD and Harmony all achieve rank 1 of 9 (*P* = 0.111), capturing the strongest attainable outcome ordering under the six-library design. This agreement, consistent with strong baseline performance in recent systematic benchmarks,^31^ shows that the frog-derived ruler captures a robust, representation-general biological contrast in mammalian repair. The mouse endpoint thus supports the reproducibility of the transported signal, while the higher-resolution frog limb comparison identifies, among the tested representations, the specific advantage of species-native tokenization over symbol collapse. Consistent with the expectation that high-dimensional steering directions need not be unique,^32^ the exhaustive source-label control—in which 4 of 10 and 3 of 10 alternative source library splits matched or exceeded the observed mouse mean—identifies the frozen ruler as a sufficient, biologically anchored direction for state transport. Together, these control hierarchies support both the cross-species generality of the repair contrast and the value of native representation in a polyploid genome.

Biologically, the competence ruler prioritized immune cells in a controlled broad-lineage screen and resolved the aligned P3-minus-P2 difference within state-balanced macrophage repair populations. Macrophages are key orchestrators of appendage regeneration in salamander limbs, zebrafish fins, and mouse digit tips,^14,33,34^ where their metabolic and activation states can influence the balance between regenerative repair and refractory fibrosis.^35,36^ Cross-species gene analysis revealed that while global transcriptomes diverge, the shared signal concentrates in a focal tissue-remodelling program involving lipid handling and matrix management (including *Apoe*, *Lpl*, *Lum*, *Psap*, *Cst3*, and *Ctsb*). This pinpoints a key cellular compartment and provides testable molecular hypotheses for tissue repair.

Prespecified external-dataset analysis in postnatal mouse skin demonstrated the developmental breadth of the competence ruler. Wounds initiated at PND2 or PND21 and collected seven days later (collected at approximately PND9 or PND28) completely separated young from old tissue. The same young-above-old ordering also appeared in unwounded skin. Because those groups were collected at different ages, this difference of contrasts is descriptive rather than an isolated injury effect. The concordant ordering across both contexts nevertheless extends the ruler-defined developmental contrast across cutaneous tissue states. Meanwhile, the reciprocal mouse-to-frog construction serves as a post hoc geometric consistency check on the forward alignment.

Deposited biological libraries, not individual cells, were the inferential units throughout. In the six-library primary mouse series, matching collection time and preserving group size leaves only nine valid outcome assignments; rank 1 of 9 (*P* = 1/9 = 0.111) is therefore the finest exact resolution available from that design. The complete 3-versus-3 library separation recurred in both UCE runs and was supported by the staged transfer sequence, depth and composition controls, state-balanced macrophage summaries and explicit null models. Additional independent libraries will sharpen the biological-label test while retaining the same analysis framework.

In summary, the tail-to-limb-to-mouse sequence establishes a framework for transporting source-defined biological contrasts as frozen, group-level rulers. Foundation embeddings thereby provide a testable way to ask whether—and in which population—a functional difference defined in one species recurs in another.

## Methods

### Study design and claim control

The forward transport chain used two frozen group-contrast rulers. The first contrasted the ROCs and ordinary epidermal cells annotated in E-MTAB-7716 and was applied unchanged to the independently embedded limb atlas. After that tail-to-limb test, the second contrasted stage-52 with stage-58 Tp63-positive limb epidermis at five days post amputation and was applied unchanged to mouse data. Mouse outcome, time, expression and annotation remained outside construction and orientation of both frog rulers.

The mouse digit analysis protocol specified datasets, matched times, sampling seeds, inferential units and success rules before download or analysis. After inspecting those results, we fixed the end-to-end reanalysis rules before balancing frog limb epidermal composition, matching mouse sequencing depth, rerunning UCE or calculating new results. This was a reanalysis of previously examined data, not preregistration. The state-balanced control-ruler analysis was specified separately before it was run. The lineage-selection and source-geometry sensitivities were specified after the primary results and before they were run. The developmental-skin analysis protocol was specified before download and preprocessing. Homeolog and ten-seed analyses are identified as post hoc or technical.

### Public single-cell datasets

The *X. laevis* tail data were obtained from ArrayExpress accession E-MTAB-7716.^27^ We retained its deposited ROC and ordinary-epidermis annotations. The independent *X. laevis* hindlimb data were obtained from E-MTAB-9104.^5^ We retained cells in its deposited “Tp63+ epidermal” group and preserved the AER annotation, developmental stage, experimental condition and sample identifier. Condition B denotes amputated limbs collected at five days post amputation; C denotes contralateral control; W denotes intact or proximal limb tissue. Supplementary Table 1 provides the complete dataset and analysis inventory.

The expression-space comparators used 7,361 marker-annotated epithelial cells from the human diabetic-foot-ulcer atlas GSE165816.^29^ These cells contributed to the 9,960-symbol cross-species feature intersection and to the unlabelled expression context jointly fitted by Harmony. Human donor outcomes remained outside ruler construction, fitting and evaluation.

The primary mouse data were obtained from GSE135985.^10^ Six deposited 10x libraries contained regenerative P3 samples at days 10 and 14 and non-regenerative P2 samples at the same times. Second-series P2 libraries at days 12, 17 and 28 were obtained from GSE293537, and matched P3 libraries from GSE267446/GSE143888.^11,13^ The primary and second dataset series were processed separately. Because all P2 and P3 samples in the second series came from different GEO studies, study and outcome cannot be separated; we therefore used these comparisons only to ask whether the P3-minus-P2 direction recurred. No new animals, human participants or tissue specimens were used.

The developmental-skin breadth data were obtained from GSE153596.^30^ The 12 deposited libraries comprise unwounded skin collected at PND2 or PND21 and wounds initiated at PND2 or PND21 and collected seven days later (approximately PND9 or PND28), with three libraries per group. The prespecified primary comparison contrasted the two 7-days-post-wounding groups by age at wounding; the unwounded developmental contrast provided context. No new animals, human participants or tissue specimens were used.

### *X. laevis* species-native protein and locus assets

Frog count-matrix features were matched to the *X. laevis* v9.1 primary-transcript annotation, primary-protein FASTA and Xenbase mappings.^28,37^ Amino-acid sequences were truncated to 1,022 residues and embedded with esm2_t48_15B_UR50D at layer 48.^26^ Following ESM2 and SATURN, final-layer representations of non-special residues were mean pooled to one 5,120-dimensional vector per protein.^19,26^ Mean pooling was retained for UCE compatibility and is not uniquely optimal.^38^ Although UCE can average multiple protein products per gene,^25^ the primary-transcript asset selected one protein per assayed feature; no across-isoform average was applied. L/S suffixes were retained, and genomic loci were obtained from v9.1 GFF3. The final asset represented 30,396 of 31,535 features; sequences, mappings and the token table are included with the reproducibility materials.

The released UCE special tokens were retained, and the *X. laevis* protein vectors were appended to the gene-token table. Species-specific chromosome tokens were initialized from a standard normal distribution using NCBI taxonomy identifier 8355 as the random seed. Only the lookup embedding changed; the released 33l_8ep_1024t_1280.torch transformer weights remained frozen.

### UCE cell inference

UCE inference used the published 33-layer transformer in evaluation mode, 1,024 expressed-gene tokens per cell sampled with replacement, padded length 1,536 and the released padding mask.^25^ Genes were sampled proportional to log(count + 1). The final classification-token (CLS) representation was L2 normalized—divided by its Euclidean length—to produce the 1,280-dimensional cell embedding.^25^ Ruler construction used the same normalization. Because released

UCE embeddings are already unit length, repeating this step does not materially alter them. Seeds 42 and 43 were used throughout. Uniform manifold approximation and projection (UMAP) served only descriptive visualization (n_neighbors=30, min_dist=0.25, cosine metric, random state 20260715), with equal x–y aspect; no UMAP coordinate entered inference.^39^

Mouse cells used the released *Mus musculus* UCE protein and chromosome vocabulary. In the primary series, 20,329 of 27,998 features were recognized; in the second series, 21,779 of a 33,529-feature union were recognized. No cell had fewer than 100 detected recognized genes. The median detected recognized-gene counts were 2,467.5 and 2,425 per cell in the primary and second series.

### Construction of frozen group-contrast rulers

For cell embedding *x*_i_, we adapted established latent difference-vector, normalized-centroid and unit-sphere formulations to a study-specific equal-library hierarchy.^21–24^ For tail library ℓ and group *g*, the construction was:

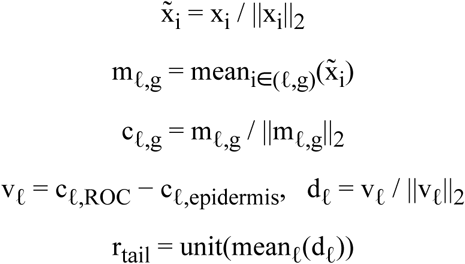

Thus, *m*_ℓ,g_ is the unrenormalized mean of unit cell vectors, *c*_ℓ,g_ is the separately unit-normalized group centroid, and *d*_ℓ_ is the unit-normalized difference used to construct the equal-library ruler. Figure 2b displays the two unit centroids after projection onto the complete ruler:

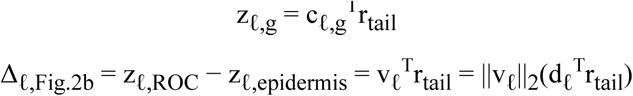

The Figure 2b connector is therefore the projection of the unrenormalized difference between two unit centroids. It is not *d* ^T^*r* itself; normalizing the difference to *d*_l_ is the subsequent step used to build the ruler. Every quantity in this panel summarizes a group centroid or the difference between two group centroids.

For computational aggregation, the code evaluated the **internal cosine coordinate** *u*_i_ = x̂ ^T^*r* for a unit ruler *r*. Because both vectors have length one, *u*_i_ is unitless and lies between −1 and +1. Its absolute value has no calibrated biological zero or threshold; it was used immediately to form prespecified within-library group summaries. A library median also lies between −1 and +1, whereas a difference between two such summaries can range from −2 to +2. Neither quantity is a probability or percentage. Standardized effects divide a raw group difference by the pooled between-library standard deviation and are therefore not bounded by either range. All biological conclusions came from contrasts between those summaries. If group means were used, mean_A_(*u*) − mean_B_(*u*) = (*m*_A_ − *m*_B_)^T^*r*, where *m* denotes the unrenormalized mean of unit cell vectors; thus, a common additive shift of the projection scalar cancels from the group difference. This identity differs from subtracting the separately unit-normalized tail-group centroids *c*. The median-based results are robust contrasts of group location.

The limb competence ruler was constructed separately within each UCE seed. Among condition-B libraries at stages 52 or 58, a unit centroid was calculated for each library. Equal-library stage centroids were formed and separately unit normalized; the normalized stage-58 centroid was then subtracted from the normalized stage-52 centroid, and the resulting difference vector was unit normalized. Target-based thresholds, fitted classifiers and sign optimization played no role in this construction.

For the controlled audit, cells annotated as AER in E-MTAB-9104 were excluded and the frog data were restricted to the five condition-B stage-52 or stage-58 libraries. The retained epidermis was divided into basal versus Znf750-positive families crossed with G1, S and G2M cell-cycle states. Unit-normalized embeddings were averaged within each of the six strata and library. Fixed stratum weights were the equal-library mean of within-library proportions; all five libraries contained at least five cells in every stratum. Weighted library centroids were unit normalized, stage centroids gave every library equal weight, and their normalized stage-52-minus-stage-58 difference defined the AER-free, composition-standardized ruler. Leave-one-library-out rulers reused the full five-library weights.

### Frog-ruler robustness and limb tests

For tail leave-one-library-out analysis, the ruler was reconstructed without each library. Robustness was measured as its cosine similarity to the complete direction. The held-out area under the receiver operating characteristic curve (AUROC) was computed from held-out ROC and epidermal cells only; libraries with fewer than five cells of either class were descriptive and excluded from the minimum eligible AUROC. The same eligible-library holdout was repeated for species-native and symbol-collapsed UCE with identical group labels, eligibility rules and seeds. Equal-library mean AUROCs were reported, and an exact one-sided paired sign test asked whether species-native UCE exceeded symbol-collapsed UCE across the ten eligible libraries.

For limb identity transfer, libraries with at least ten AER and ten other Tp63-positive cells were summarized as median internal cosine coordinate in AER minus median internal cosine coordinate in other epidermis. The exact one-sided sign-flip test enumerated all 2*^n^* sign assignments.

For developmental competence, each condition-B library was summarized by the median internal cosine coordinate across its epidermal cells. The primary statistic was the equal-library mean of the three stage-52 summaries minus the equal-library mean of the two stage-58 summaries; stage 56 remained in the label-enumeration design but not in this contrast. Exact inference enumerated all stage-label allocations across the eight condition-B libraries while preserving the 3/3/2 stage-52/56/58 sizes. Standardized effects divided this raw contrast by the pooled library-level standard deviation and are descriptive because only three stage-52 and two stage-58 libraries were available.

For AER-abundance sensitivity, annotated AER cells were removed before library summarization and stage-label enumeration was repeated. This tests whether organizer abundance alone accounts for the contrast; it does not balance all remaining epidermal substates.

### Token-sampling robustness audit

The canonical UCE sampling procedure was repeated for seeds 44–51 in addition to the two prespecified seeds 42 and 43, yielding ten independently sampled embeddings for tail and limb cells. Rulers were constructed for every tail seed and applied to every limb seed (100 combinations). The audit recorded raw stage-52-minus-stage-58 contrasts, complete early/late separation, stage-group median order and exact stage-label probabilities. Ten-seed ensembles were formed by averaging unit-length cell embeddings across seeds and scaling the average back to unit length; each seed was then omitted in turn to assess convergence. Alternative gene-selection procedures were exploratory and are not part of the primary analysis.

### Homeolog-resolved expression audit

For each gene in the homeolog-resolved limb count matrix, equal-library stage-52-minus-stage-58 log-normalized expression effects were calculated among condition-B epidermal cells. Base symbols with both an L and S copy yielded 7,702 pairs. Pair-level Spearman correlation, same-direction fractions and the high-effect subset were calculated across all pairs; high-effect pairs had at least one copy above the genome-wide 90th percentile of absolute single-gene effect. Pair correlation and focal copy-asymmetry classes were recalculated after omitting each of the five stage-52 or stage-58 libraries. The analysis is descriptive and post hoc.

### Symbol-collapsed UCE and expression-space comparisons

For symbol-collapsed UCE, *X. laevis* features were mapped to *X. tropicalis* symbols present in the released UCE protein and chromosome vocabularies; duplicate mappings were summed. The frozen transformer and seeds were unchanged. The expression-space comparisons used 9,960 uppercase symbols shared by the frog matrices and GSE165816 human epithelium after removing .L/.S suffixes and summing homeolog counts. Counts were normalized to 10,000, log1p transformed and unit normalized. Direct expression used this feature space without a fitted reduction. Tail-trained SVD fitted 128 components on tail cells and then applied the transformation unchanged to limb cells. Harmony fitted 50 joint components from tail, limb and human epithelial expression and corrected dataset identity; it used target expression but no AER, stage or human outcome labels.^16,29^ These tests compare the integrated representation-and-ruler routes; they do not isolate one architectural component of UCE.

### Mouse quality control, blind annotation and cell sampling

Mouse cells were retained when they contained 200–6,500 detected genes, at least 500 total counts and no more than 20% mitochondrial counts. The primary series contained 11,344 cells before and 10,433 after filtering; second-series library counts after filtering ranged from 3,351 to 13,497. Outcome and time columns were carried in a separate evaluation table and were not provided to UCE, clustering or subsampling.

For broad lineage annotation, each dataset series was normalized to 10,000 counts per cell and log transformed. Three thousand highly variable genes and 50 principal components were calculated with Scanpy.^40^ K-means clustering with 16 clusters, random state 135985 and 20 initializations was performed without outcome or time. Cluster means were calculated for predeclared marker panels: mesenchymal (*Col1a1, Col1a2, Col3a1, Dcn, Lum, Pdgfra, Col5a1, Col6a1*); epidermal (*Krt5, Krt14, Krt15, Krt17, Krt19, Krt1, Krt10*); endothelial (*Pecam1, Cdh5, Kdr, Emcn, Esam, Klf2*); immune (*Ptprc, Lyz2, Csf1r, Tyrobp, Cd68, Spi1*); mural (*Rgs5, Pdgfrb, Cspg4, Acta2, Tagln, Myh11*); neural (*Sox10, S100b, Plp1, Mpz, Sox2, Fabp7*); osteogenic (*Sp7, Runx2, Bglap, Ibsp, Spp1, Alpl*); and erythroid (*Hba-a1, Hba-a2, Hbb-bs, Hbb-bt, Alas2, Gypa*). Each cluster was assigned to the panel with the highest score standardized by the within-panel median and median absolute deviation (MAD).

To equalize compute and prevent the largest library from dominating, the primary series was sampled to 692 cells per library while approximately preserving its blind-cluster composition. Second-series libraries were sampled randomly to 1,000 cells each. Selection used fixed seeds and no outcome/time values.

### Mouse outcome tests

For the initial tissue-level screen, each deposited library was summarized by the median internal cosine coordinate obtained with the corresponding seed-specific frog competence ruler. Biological interpretation used the difference between condition-level library summaries. Within each primary-series day, we calculated the mean P3 library summary minus the mean P2 library summary; the time-adjusted raw contrast was the unweighted mean of the day-10 and day-14 differences. The exact randomization distribution enumerated every outcome allocation within day while preserving P3 library counts. Because only nine allocations exist, the minimum attainable one-sided probability is 1/9.

The second-series screen used the difference between the P3 and P2 library summaries at days 12, 17 and 28. The two dataset series were summarized separately. Direction across the two primary and three second-series times is temporal consistency only: repeated times within a series are not independent biological experiments, and outcome is aligned with accession in the second series. A random-series mixed model was not fitted because two levels are insufficient for reliable estimation of between-series variance.

### Pre-inference depth matching and whole-tissue scope audit

Every retained mouse cell was randomly downsampled to exactly 500 raw UMI counts by multivariate hypergeometric sampling without replacement. Cell-specific seeds derived with the BLAKE2b hash algorithm made the draw independent of row order. The operation retained all 4,152 sampled primary cells and all 6,000 sampled second-series cells. UCE was rerun from these count matrices with the published checkpoint, mouse vocabulary, 1,024-gene sample, 1,536-token pad length and seeds 42/43 unchanged. No post-inference depth regression was used. Embeddings were scaled to unit length and compared with the seed-matched AER-free, composition-standardized frog ruler.

Whole-tissue audit summaries used deposited-library medians of the internal cosine coordinates and the same five matched P3-minus-P2 contrasts as the initial screen. The primary statistic and its nine blocked outcome allocations were unchanged. The second-series contrasts mapped the reach of whole-tissue consistency across studies.

### Developmental skin and hair-follicle breadth test

GSE153596 cells were retained when they contained 200–6,500 detected genes, at least 500 total counts and no more than 20% mitochondrial counts. This left 66,921 cells. Condition labels were held outside preprocessing. Sixteen blind clusters were assigned to broad lineages from prespecified marker panels, and 700 cells per deposited library were selected by proportional sampling within blind clusters, yielding 8,400 embedded cells. Mouse UCE inference used the same frozen checkpoint, frog competence rulers and prespecified seeds 42 and 43 used for the digit analysis.

Each skin library was summarized by its median internal cosine coordinate. The primary statistic was the equal-library mean for samples wounded at PND2 minus the corresponding mean for samples wounded at PND21; both groups were collected seven days after wounding. Exact inference enumerated all 20 three-versus-three allocations (minimum *P* = 0.05). The unwounded PND2-minus-PND21 contrast used the same calculation. Because the wounded and unwounded groups were collected at different absolute ages, the difference between these contrasts was retained as a descriptive quantity and was not interpreted as an isolated injury interaction.

Composition robustness imposed equal broad-lineage contributions, requiring at least 20 cells per library-lineage stratum. Sampling robustness repeated equal selection of 300 cells per library 1,000 times per seed. Matched shared-gene expression and tail-trained SVD rulers used 11,584 shared symbols and the same exact test. These analyses quantify developmental breadth across skin contexts.

### Depth-matched lineage and macrophage-state analyses

The broad-lineage screen used the depth-matched mouse embeddings and outcome-blind annotations. A lineage was eligible for the complete screen when both outcomes contributed at least 20 cells at each of the five matched comparisons. Within each lineage, every deposited library was summarized by its median internal cosine coordinate, and each matched contrast was the mean P3 summary minus the mean P2 summary. The immune compartment was selected after examining these outcome contrasts, so the lineage-selection step is outcome-informed.

Within depth-matched immune cells, counts were transformed as log1p(10,000 × count / 500) and each gene was standardized within dataset series. Macrophage-like cells had a macrophage-identity module (*C1qa, C1qb, Adgre1, Mrc1, Csf1r, Cd68*) above a neutrophil module (*S100a8, S100a9, Retnlg, Ly6g, Mmp9*). Four fixed state modules were then scored: lipid/phagocytic (*Apoe, C1qa, C1qb, C1qc, Trem2, Lpl*), resident/remodelling (*Mrc1, Folr2, Lyve1, Cd163, Cbr2*), inflammatory (*Il1b, S100a8, S100a9, Cxcl2, Ptgs2*) and interferon (*Isg15, Ifit1, Ifit3, Irf7, Oasl2*). These names are operational, literature-grounded program labels rather than claims of discrete universal macrophage ontologies.^41–43^ Each cell was assigned to the module with the highest within-library robust z score, calculated from the median and MAD, with ties resolved in the fixed module order. Modules required at least three present genes.

A state was eligible separately within each series only if every deposited library contained at least ten assigned cells; at least two states were required and no fallback was allowed. Within each eligible state, the cell-level internal cosine coordinates were reduced to a median; each library summary was then the equal mean of these state medians. This yielded two eligible states in the primary series and three in the second series. At each matched time, the reported contrast was the mean P3 library summary minus the mean P2 library summary. Primary exact probabilities enumerated the same nine outcome-label assignments while preserving matched day; the five-sign probability enumerated all 2^5^ sign assignments and was treated as corroborating.

For leave-one-library-out analysis, the AER-free ruler was reconstructed after omitting each of five frog libraries while retaining the full-data composition weights. All ten allocations of three stage-52 and two stage-58 labels across those libraries were enumerated. The biological control ruler contrasted cycling and G1 frog epidermal cells with the same family balancing and equal-library weighting. The original geometric null comprised 10,000 random unit directions sampled without a preferred orientation from a Gaussian distribution per seed. All controls used identical mouse cells, eligible states and aggregation; the macrophage test statistic was the unweighted mean of five P3-minus-P2 contrasts.

A post hoc adjustment accounted for broad-lineage selection. For each sampled direction, eligible lineages were ranked first by the number of positive matched contrasts and then by the mean of the five contrasts. A draw met or exceeded the observed immune result only when the selected lineage had five positive contrasts and a mean at least as large as the immune-lineage mean. This rule was applied to 10,000 uniformly sampled full-space directions per UCE run.

A second post hoc null sampled directions from frog source-supported geometry. Thirty unit centroids—six AER-free epidermal-family-by-cell-cycle strata in each of five frog libraries—were centered and decomposed by principal components without mouse labels. The first five components retained at least 90% of source variance in each UCE run. Gaussian coefficients were scaled by the corresponding source singular values, combined across those five components and normalized to unit length. Ten thousand such directions per run were evaluated with both the repeated lineage-selection rule and the fixed state-balanced macrophage summary. This covariance-matched null tests sensitivity to anisotropic source geometry and does not replace the exhaustive ten-allocation source-label control. Specifications, code and complete output tables are provided with the reproducibility materials.

Sampled-null probabilities used (1 + number null ≥ observed) / (number sampled + 1);^44^ exhaustive allocations were reported as exact fractions. Tail ROC labels were permuted 1,000 times within library while preserving class counts, and each resulting ruler was evaluated on limb AER separation and the composition-standardized non-AER stage contrast. State-balanced cross-application and the selection-aware controls were specified after the core analysis.

### Controlled mouse-to-frog reciprocity stress test

After completing the forward analysis, we constructed a post hoc reverse ruler from the six depth-matched GSE135985 primary mouse libraries. Within each library, unit-length cell embeddings were averaged separately for the lipid/phagocytic and inflammatory macrophage states that met the fixed eligibility rule in every library. The two state means contributed equally, and their average was rescaled to unit length to define one library centroid. Dimension-wise least squares across the six centroids included an intercept, a day-14 indicator and a regenerative-P3 indicator; the unit-normalized P3 coefficient defined the frozen mouse ruler.

The mouse ruler was applied without refitting to the frog limb and tail embeddings. Each condition-B limb library was summarized by its median internal coordinate, and the stage-52-minus-stage-58 statistic used all 560 allocations preserving the observed 3/3/2 stage counts. The AER-free sensitivity reused the fixed six-stratum epidermal weights and enumerated all ten three-versus-two stage allocations. Tail libraries were summarized by the median ROC coordinate minus the median ordinary-epidermis coordinate, with exact inference over all 2^14^ sign flips. Robustness refitted the ruler after each of six mouse-library omissions and under all nine within-day mouse outcome allocations. Complete results and code are provided with the reverse-transfer materials.

### Orthology-restricted molecular-bridge decomposition

Shared uppercase gene symbols were defined across frog limb, primary mouse and second-series mouse matrices after removing *X. laevis* .L and .S suffixes and summing homeolog counts. Each matrix was library-size normalized to 10,000 and log1p transformed. For frog, the gene effect was the difference between equal-library mean expression at stage 52 and stage 58 among condition-B epidermal cells. For each mouse dataset series, library-level P3-minus-P2 expression effects were calculated within the immune lineage and averaged across matched times. Spearman correlations used all 11,549 shared symbols. The focused display includes selected high-ranking non-translation genes positive in all three contrasts together with the previously defined immune marker set.

The repair/lipid/phagocyte set was tested for over-representation among genes positive in all three contrasts. It comprises the lipid/phagocytic module plus *Mrc1*; the acute-inflammatory set matches the inflammatory module, so marker interpretation and state naming are not independent. Bridge effects were calculated across the whole immune lineage without conditioning on state assignment. Genes absent from the 11,549-symbol universe were excluded.

A one-sided exact hypergeometric test fixed the gene universe, set size and number of genes positive in all three contrasts. Fold enrichment was observed divided by expectation; correction covered two directional set tests. This gene-label test is distinct from replicate inference. The decomposition is post hoc and interprets the output of the ruler rather than attributing individual UCE dimensions: protein tokens undergo nonlinear encoding and 33 transformer layers before application of the 1,280-dimensional direction.

### End-to-end representation comparison

To establish whether the primary mouse endpoint discriminates between representations, the complete forward chain was rerun independently for direct expression, source-only SVD and Harmony alongside both species-native UCE sampling runs. Each representation built its own unit-normalized stage-52-minus-stage-58 frog ruler from the same five AER-free limb libraries after balancing basal and Znf750 epidermal families and G1/S/G2M states, then applied that ruler without outcome-label fitting to the same 4,152 mouse cells rarefied to 500 raw UMIs. Expression-space comparators used the 11,682 gene symbols shared between the frog and mouse feature spaces. Harmony was fitted jointly on source and target expression but remained blind to mouse outcome labels; direct expression and SVD were fitted on source only. Cell coordinates were summarized by deposited library, and all nine within-day outcome allocations were enumerated to give an exact one-sided probability. The comparison was specified after the primary mouse result had been seen and is therefore a post hoc audit on previously examined data, not a preregistered test. Complete results and code are provided with the reproducibility materials.

### Statistics, blinding and reporting

Deposited libraries—not cells—were the inferential units for biological contrasts.^45^ Exact enumeration was used where feasible and preserved experimental matching. Five mouse times nested within two series were not treated as independent cohorts. Cell-level correlations, UMAPs, standardized effects and gene-level tests are labelled descriptive, technical or interpretive.

Mouse outcome, time, age and wound labels were withheld from embedding, blind clustering and subsampling and restored for frozen-ruler evaluation. The end-to-end audit rules were fixed after inspecting the mouse data; the selection-aware and source-geometry controls were specified after completing the core analysis. All eligible libraries were reanalysed, so no power calculation was performed. No cell or library was excluded after directional results were inspected beyond frozen quality-control, frog-stratum, lineage-completeness and state-eligibility rules.

### Reproducibility and software

All processing, model-inference, analysis and figure-rendering scripts are organized within the accompanying reproducibility materials. Major packages were Python, PyTorch, Scanpy, anndata, NumPy, pandas, SciPy, scikit-learn, harmonypy, UMAP-learn, Matplotlib and Seaborn. These materials include executable scripts, exact result tables, ruler vectors and provenance records; large depth-matched matrices and regenerable embeddings are listed in the external-data manifest. The developmental-skin, reverse-transfer and selection-aware control analyses are provided as separate modules. Public raw matrices, model checkpoints and other large artifacts are also enumerated in the external-data manifest. No UCE model weight was altered.

## Data availability

All primary data are publicly available from ArrayExpress (E-MTAB-7716 and E-MTAB-9104) and the Gene Expression Omnibus (GEO; GSE165816, GSE135985, GSE293537, GSE267446, GSE143888 and GSE153596). The reproducibility materials contain frozen ruler vectors, intermediate cell-level internal cosine coordinates, library- and group-level results, figure source data, scripts and provenance manifests. Public raw-data locations and omitted regenerable large artifacts are listed in the external-data manifest. The associated analysis and figure-rendering code is available through the public repository described below.

## Code availability

All code needed to reconstruct the analyses and figures is publicly available at https://github.com/hucang0/uce-competence-ruler. The code-only repository provides a public-data manifest, named preparation and analysis entry points, the checkpoint-compatible UCE inference implementation and renderers for every main and supplementary figure. Datasets, model checkpoints, generated embeddings, result tables and images are not redistributed in the repository; its dataset manifest records the public accessions and upstream asset locations needed to regenerate them locally. The published UCE checkpoint and released species assets remain governed by the original UCE distribution terms.^25^

## Ethics statement

This study is a computational reanalysis of publicly available de-identified data and did not involve new animal experiments or human participants.

---

**Supplementary Figure 1.**
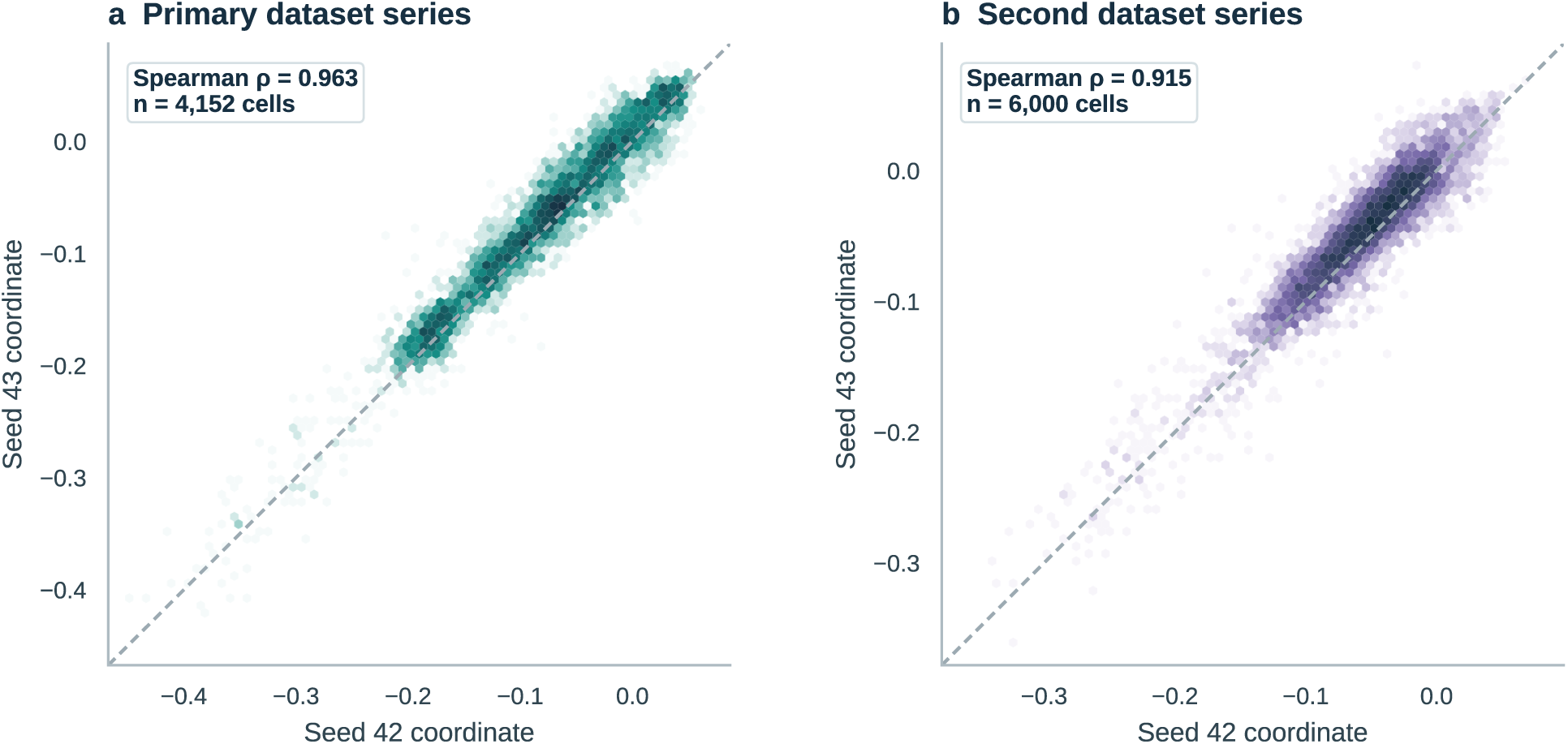
Cell-level token-sampling robustness in mouse. Each axis is the intermediate internal cosine coordinate—the dot product of a unit cell vector and its seed-matched unit ruler, bounded from −1 to +1—for seed 42 or 43 in the primary and second dataset series. These same-cell correlations assess computational reproducibility; biological conclusions compare deposited-library and condition-group summaries.

**Supplementary Figure 2.**
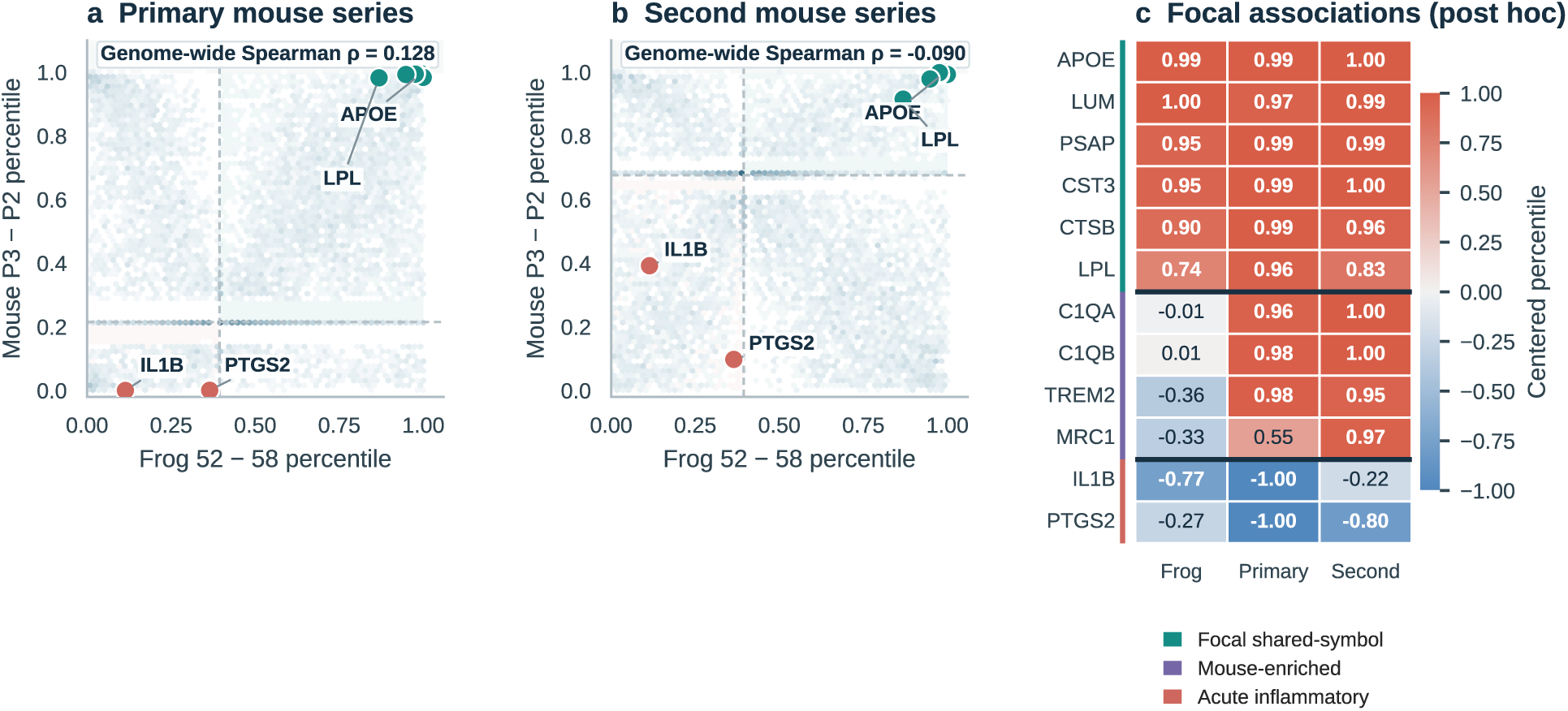
A focal frog-to-mouse tissue-remodelling bridge. a,b, Percentile-ranked shared-symbol expression effects for frog stage-52 versus stage-58 epidermis and immune-lineage P3 versus P2 effects in the primary and second mouse dataset series. Grey points show all 11,549 shared symbols and shaded quadrants show sign concordance; teal points mark selected non-translation genes positive in all three contrasts. **c,** Centered within-contrast percentiles distinguish focal lipid/lysosomal/extracellular-matrix-remodelling candidates from the mouse-dominant complement/*Trem2/Mrc1* component. The repair-module enrichment is reported in Supplementary Table 3. This orthology-restricted, post hoc analysis provides molecular interpretation of the transported ruler.

**Supplementary Figure 3.**
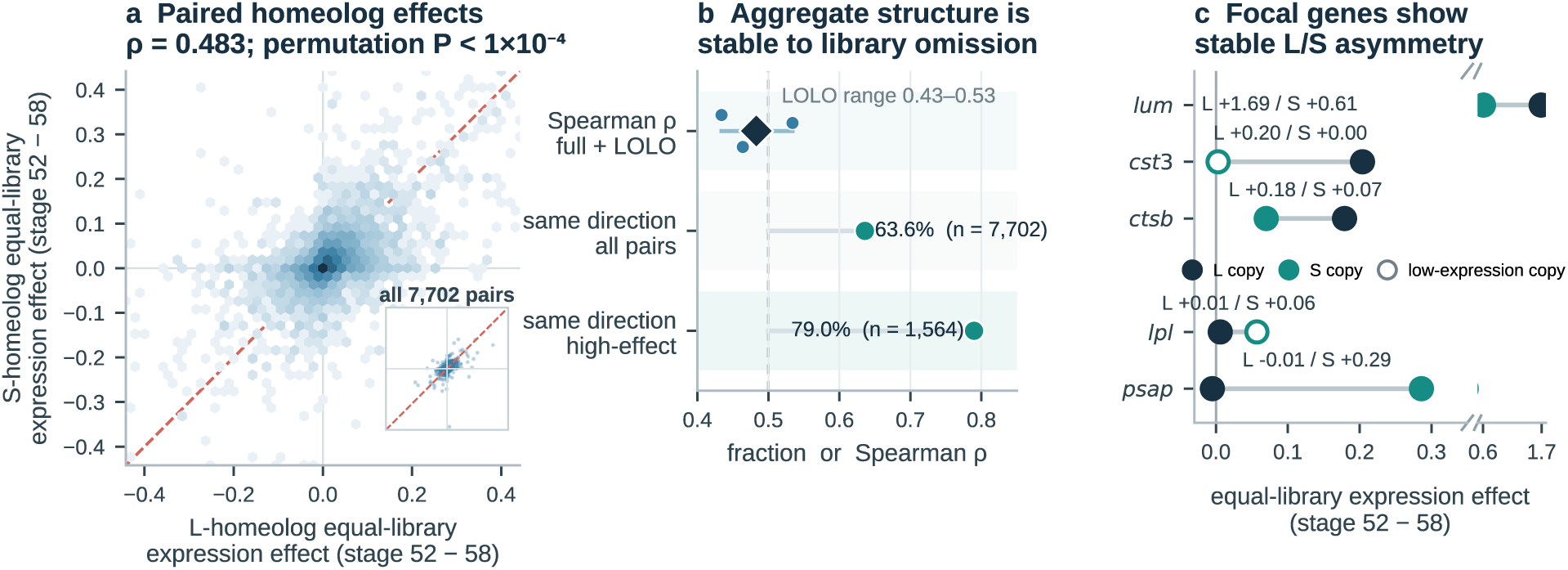
Homeolog-resolved expression structure in the frog limb competence contrast. a, Equal-library stage-52-minus-stage-58 log-normalized expression effects for 7,702 paired *Xenopus laevis* L/S homeologs; all pairs enter the statistics (Spearman ρ = 0.483; 10,000-permutation *P* < 1 × 10^−4^). **b,** Same-direction fractions across all and high-effect pairs, with full-data and leave-one-library-out correlations. **c,** L- and S-copy effects for focal genes whose asymmetry class was retained across all five library omissions; open symbols mark low-expression copies. This post hoc audit characterizes the homeolog structure retained by species-native UCE.

**Supplementary Figure 4.**
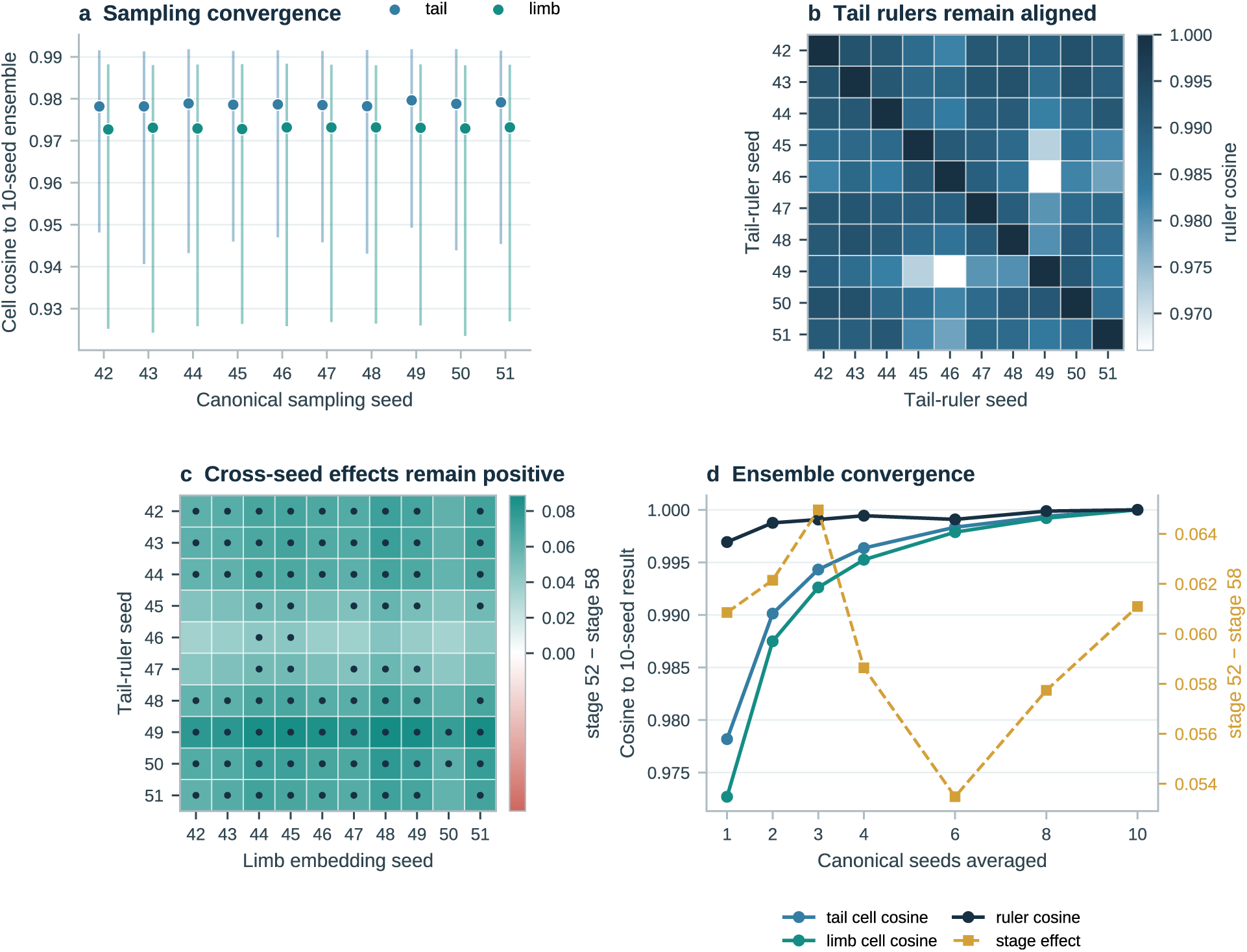
Ten-seed robustness of the frog tail-to-limb transfer. Canonical UCE token sampling was repeated for seeds 42–51. **a,** Same-cell cosine similarities compare each seed-specific UCE embedding vector with the unit-normalized ten-seed mean vector. **b,** Pairwise cosine similarities among the ten tail rulers. **c,** All 100 tail-seed-by-limb-seed combinations produced a positive raw stage-52-minus-stage-58 coordinate difference and complete early/late library separation; dots mark combinations with exact *P* ≤ 0.05 and the expected median ordering. **d,** Convergence of same-cell embedding cosines, ruler cosine and the raw stage coordinate difference as seeds enter the ensemble. The ten-seed ensemble raw coordinate difference was 0.0611 with exact *P* = 0.0339. All ten leave-one-seed-out ensembles preserved direction, exact threshold and complete stage-52/58 separation. The mouse ruler remains based on the two prespecified seeds.

**Supplementary Figure 5.**
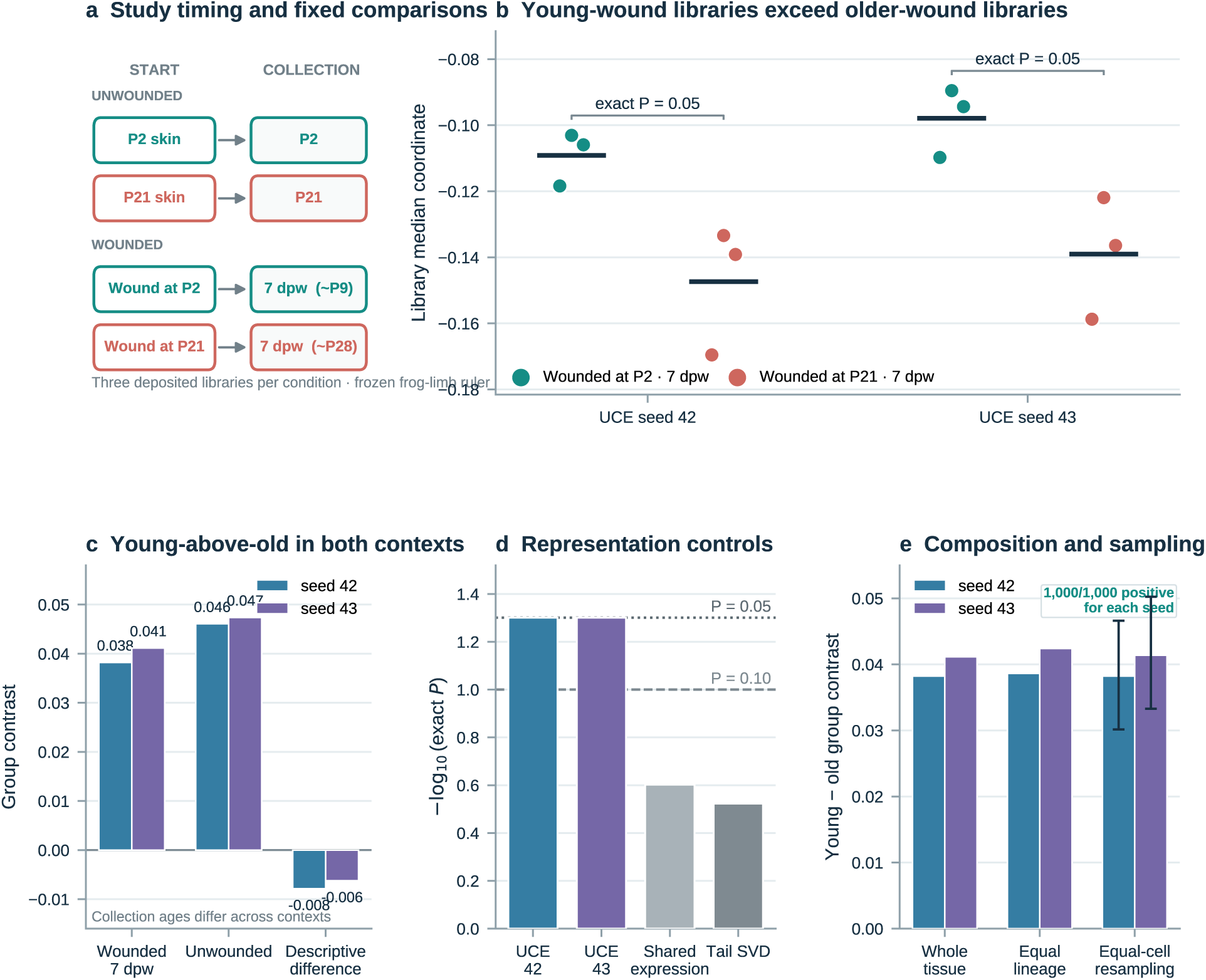
A frozen frog-limb ruler detects a developmental contrast across uninjured and wounded skin. a, GSE153596 timing. Unwounded skin was collected at PND2 or PND21; wounds initiated at PND2 or PND21 were collected seven days later (approximately PND9 or PND28), with three deposited libraries per group. **b,** Each point is a library median internal cosine coordinate; all libraries wounded at PND2 exceed all libraries wounded at PND21 for both UCE seeds, bars denote equal-library means and the exact one-sided probability is 0.05. **c,** Young-minus-old contrasts in wounded and unwounded contexts; their difference is descriptive because collection ages differ. **d,** Matched comparisons across 11,584 shared symbols place the UCE result against shared-gene expression and tail-trained SVD rulers. **e,** The wounded contrast remains positive after equal-lineage reweighting and in all 1,000 equal-cell resamples per seed. Raw contrasts are unitless coordinate differences, not probabilities.

**Supplementary Figure 6.**
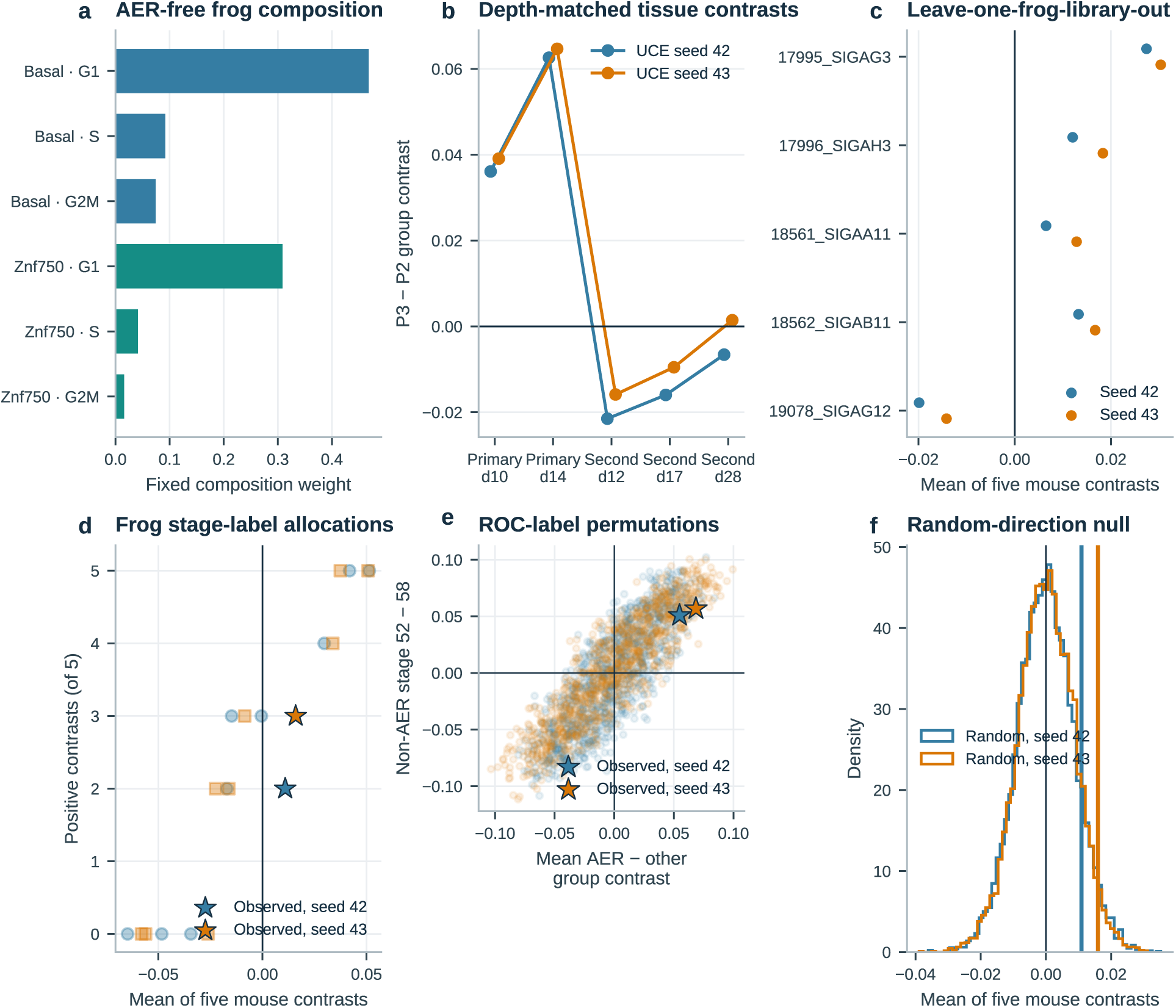
Controlled analyses resolve tissue-level transfer and upstream label evidence. a, Fixed weights across six AER-free frog epidermal strata. **b,** Pre-inference depth-matched tissue contrasts. **c,** Means of the five whole-tissue raw contrasts after leaving out each frog library. **d,** All ten frog-stage label allocations evaluated on the same five-contrast mean. **e,** Tail ROC labels permuted within each E-MTAB-7716 library; stars mark the observed AER separation and composition-standardized non-AER developmental contrast. **f,** The observed five-contrast whole-tissue mean against 10,000 random directions. Together, these controls motivate the progression from whole-tissue search to state-balanced macrophage analysis.

**Supplementary Figure 7.**
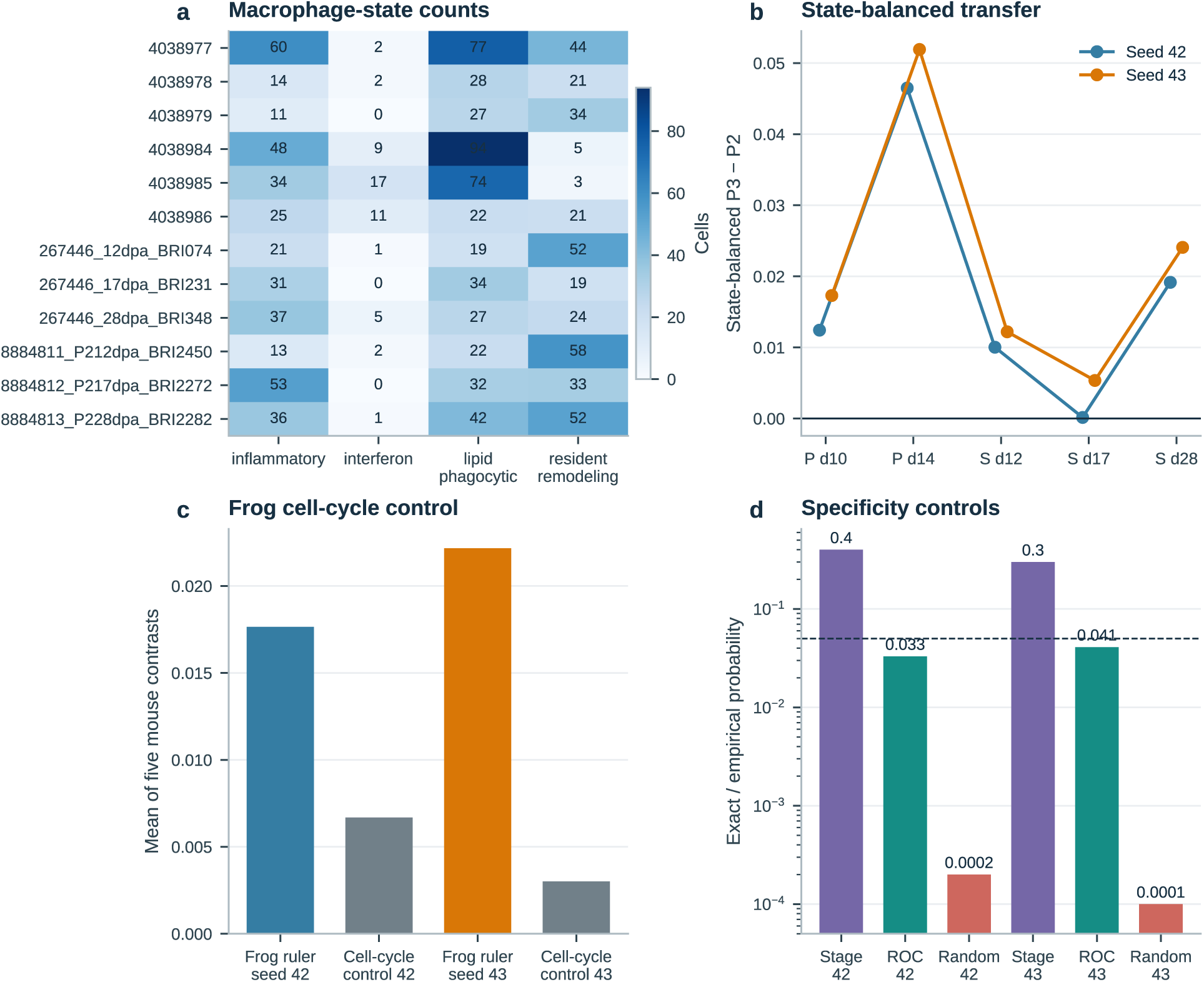
Fixed macrophage-state eligibility and summarized controls. a, Cell counts in four fixed macrophage states for every deposited mouse library; eligibility required at least ten cells in every library of a series. **b,** State-balanced P3-minus-P2 contrasts. **c,** Mean of the five state-balanced mouse contrasts for the frog competence ruler and frog cell-cycle control ruler. **d,** Exact frog-stage, joint ROC-permutation and random-direction probabilities on a logarithmic axis. The heat map displays every state count, including ineligible interferon and primary resident/remodelling strata.

**Supplementary Figure 8.**
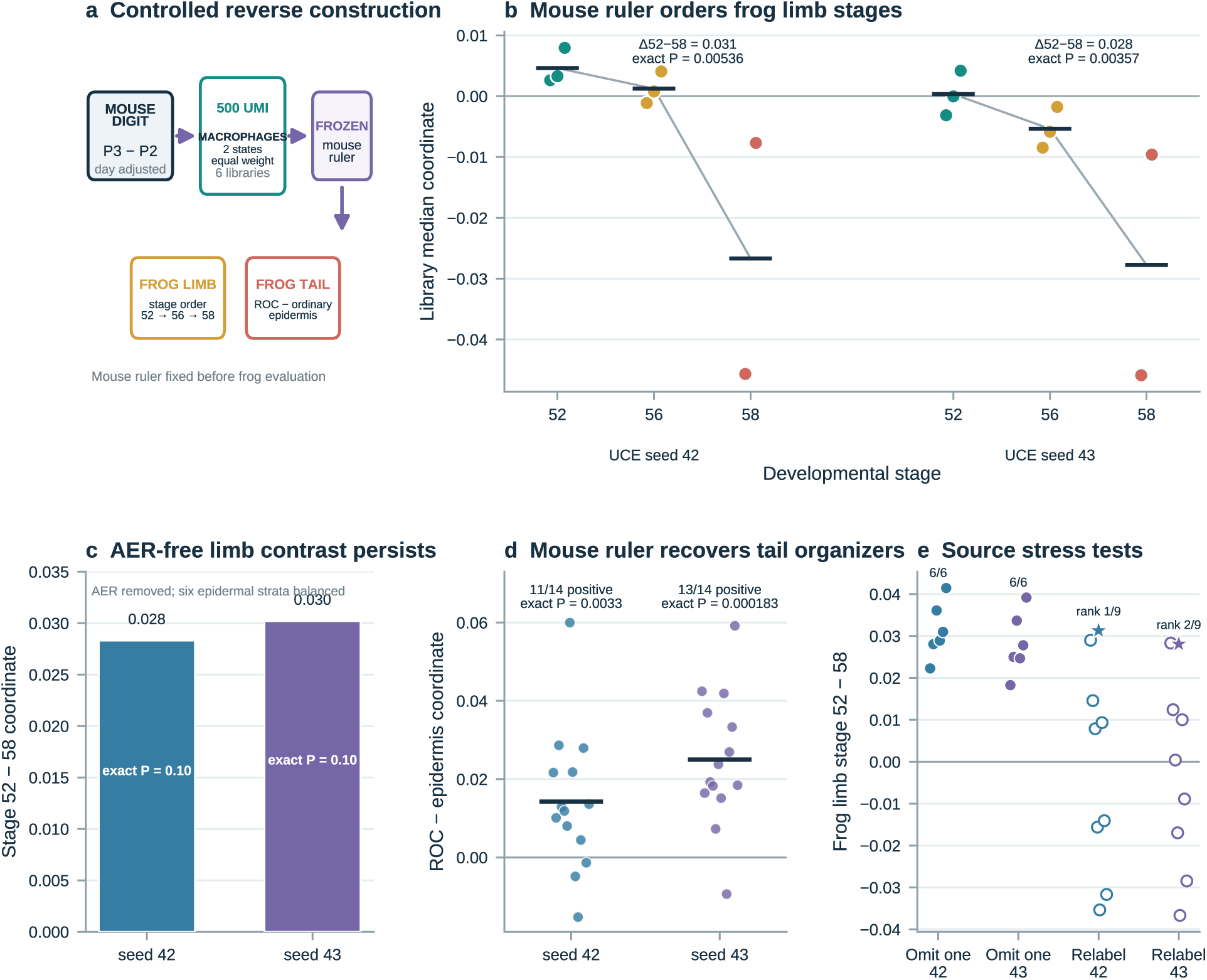
Reciprocal mouse-to-frog construction provides a geometric consistency check. a, The reverse ruler was built from six 500-UMI primary mouse digit libraries after giving lipid/phagocytic and inflammatory macrophage states equal weight and adjusting the P3-versus-P2 direction for collection day; frog data entered only after the ruler was frozen. **b,** Deposited-library median coordinates in the independent frog limb atlas. Both UCE runs produced the strict stage-52 > stage-56 > stage-58 order; horizontal bars denote equal-library stage means. **c,** Stage-52-minus-stage-58 differences after AER removal and six-stratum epidermal balancing. **d,** Within-library tail ROC-minus-ordinary-epidermis differences; horizontal bars denote equal-library means. **e,** Frog limb stage gaps after each mouse source-library omission (filled points) and across all nine within-day mouse outcome-label allocations (open points); stars mark the observed labels. The mouse-derived ruler was reconstructed independently for each UCE sampling run.

**Supplementary Figure 9.**
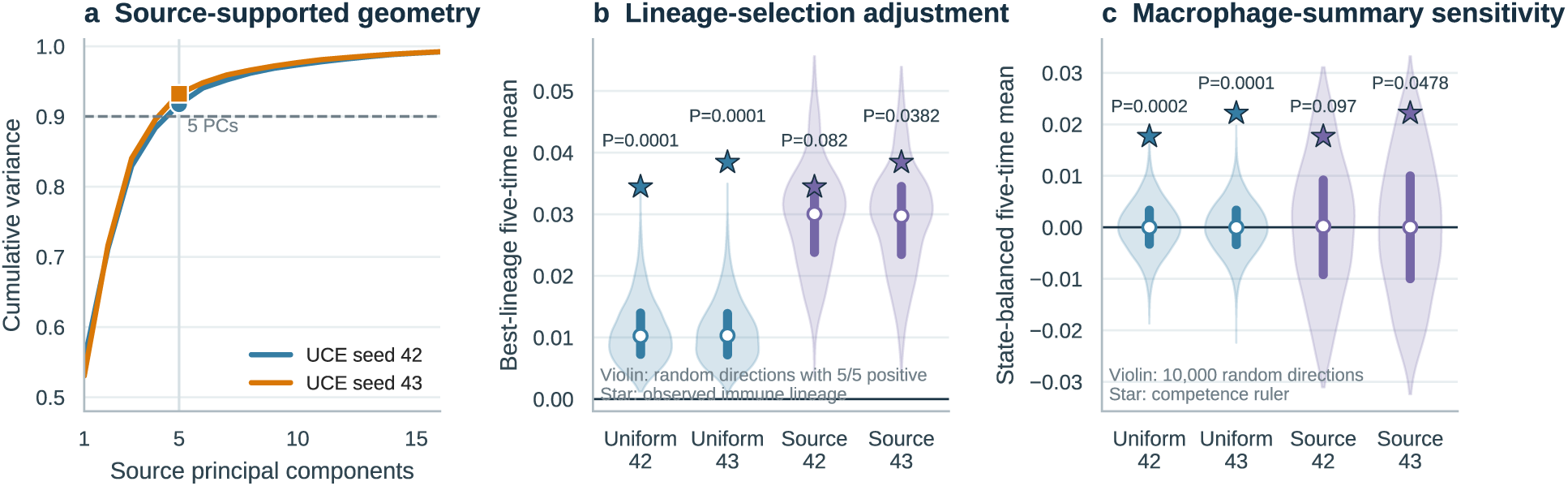
Selection-aware and source-supported controls calibrate ruler specificity. a, Cumulative variance of 30 AER-free frog stratum-by-library centroids; five principal components retain 91.7% and 93.2% of source variation in the two UCE runs. **b,** For every random direction, the complete lineage with the most positive P3-minus-P2 contrasts and then the largest five-time mean was selected. Stars mark the observed immune-lineage result; violins show uniformly sampled full-space directions or directions matched to frog source covariance that achieved five positive contrasts. **c,** The same direction families evaluated with the state-balanced macrophage summary fixed in advance. Each null contains 10,000 directions per UCE run, and displayed probabilities use the +1 correction.

**Supplementary Figure 10.**
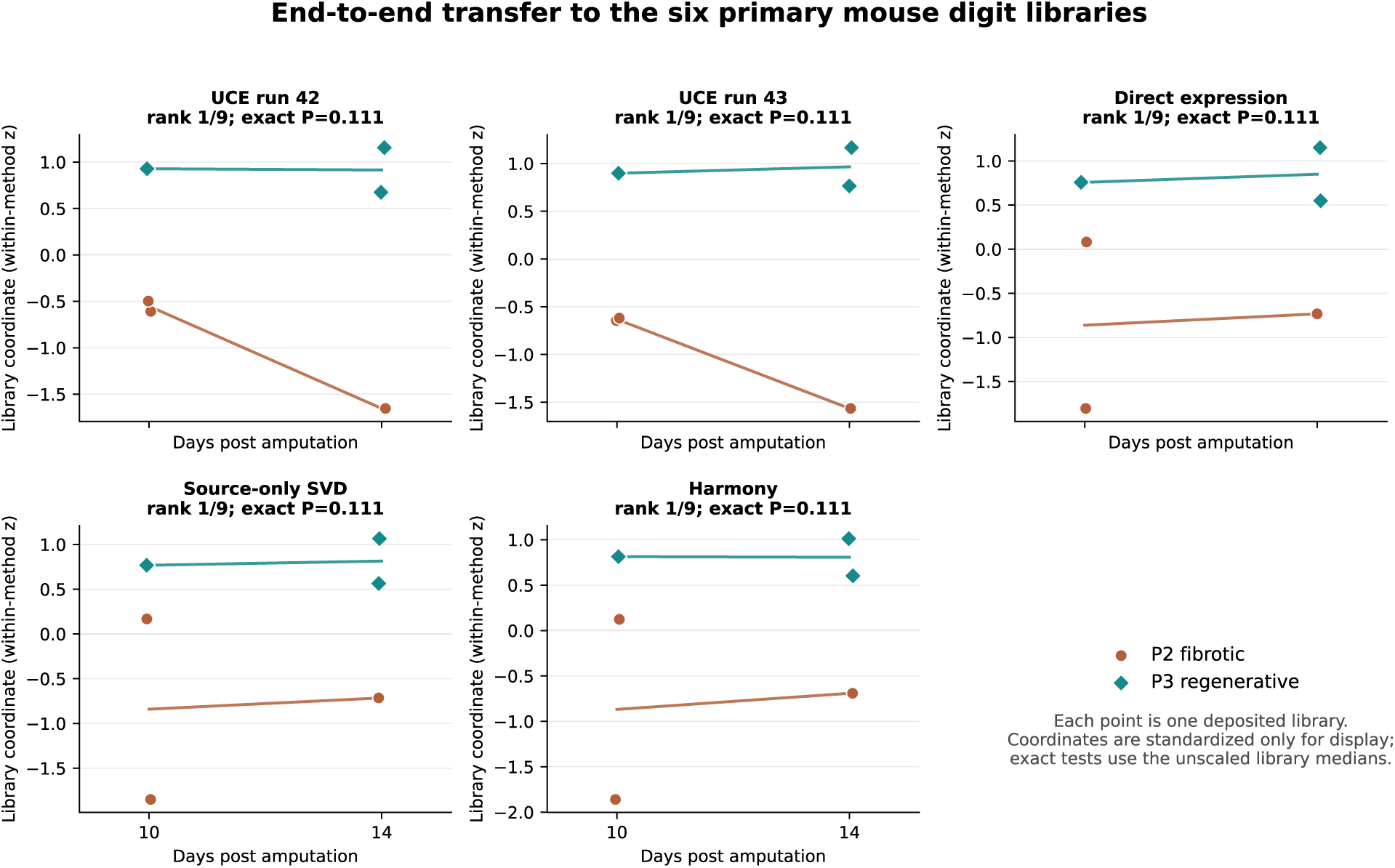
The primary mouse endpoint is reproduced by simpler representations. The complete frog-to-mouse chain was rerun independently for each representation. Every representation constructed its own frozen stage-52-minus-stage-58 ruler from the same five AER-free frog limb libraries, balanced across basal and Znf750 epidermal families and G1/S/G2M states, and applied it without outcome-label fitting to the same 4,152 mouse cells rarefied to 500 UMIs. Each point is one deposited library summarized by its median internal coordinate; coordinates are standardized within representation for display only, and the exact tests use unscaled library medians. All nine within-day outcome allocations were enumerated for each representation. Species-native UCE (both sampling runs), direct expression, source-only SVD and Harmony each place the observed labels first of nine, give complete 3-versus-3 separation and remain positive in all five source leave-one-out omissions. Harmony is target-aware but outcome-blind; direct expression and SVD are source-only. Raw coordinate magnitudes are not comparable across representations. Numerical values are given in Supplementary Table 4, and complete results and code are provided with the reproducibility materials.

**Supplementary Table 1.** Dataset and analysis inventory.

| Role | Accession | Species / tissue | Cells used here | Deposited libraries / donors | Biological contrast |
| --- | --- | --- | --- | --- | --- |
| Source-ruler construction | E-MTAB-7716 | *X. laevis* tail epidermis | 2,054 | 14 | ROC versus ordinary epidermis |
| Independent limb-transfer test | E-MTAB-9104 | *X. laevis* limb<br>Tp63+ epidermis | 9,142 | 18 | AER identity; stage 52 versus 58 at 5 dpa |
| Expression-comparator context | GSE165816 | *H. sapiens* diabetic-foot-ulcer epithelium | 7,361 | 11 donors | 9,960-symbol intersection and unlabelled Harmony context; outcomes unused |
| Mammalian primary series | GSE135985 | *M. musculus* digit injury | 4,152 | 6 | regenerative P3 versus fibrotic P2 at 10 and 14 dpa |
| Mammalian second series | GSE293537 /<br>GSE267446 /<br>GSE143888 | *M. musculus* digit injury | 6,000 | 6 | P3 versus P2 at 12, 17 and 28 dpa |
| Developmental skin / hair-follicle breadth test | GSE153596 | *M. musculus* skin / hair follicle | 8,400 sampled (66,921 after quality control) | 12 | young versus old skin; wounds initiated at PND2 or PND21 and collected 7 days later |

*Abbreviations.* AER, apical ectodermal ridge; dpa, days post amputation; PND, postnatal day; ROC, regeneration-organizing cell.

**Supplementary Table 2.** Principal results and controlled mouse analyses.

| Comparison or test | Seed 42 | Seed 43 | Inferential unit |
| --- | --- | --- | --- |
| Tail minimum leave-one-library-out direction cosine | 0.9973 | 0.9975 | held-out source library |
| Tail minimum eligible held-out AUROC | 0.8728 | 0.8377 | held-out source library |
| Limb AER-positive libraries | 12 / 14 | 12 / 14 | deposited library |
| Limb AER exact one-sided probability | 0.000549 | 0.000244 | library sign flip |
| Limb stage 52 versus 58 descriptive standardized effect | 4.226 | 6.519 | 3 stage-52 versus 2 stage-58 libraries |
| Limb stage 52 minus 58 raw coordinate contrast | 0.06085 | 0.06308 | equal-library means; 3 stage-52 versus 2 stage-58 libraries |
| Limb exact stage-label probability | 0.0393 | 0.0250 | deposited 5-dpa library |
| Non-AER limb stage 52 minus 58 raw contrast | 0.0391 | 0.0385 | post hoc AER-exclusion sensitivity |
| Non-AER exact stage-label probability | 0.0929 | 0.0786 | deposited 5-dpa library |
| Initial mouse tissue screen: primary time-adjusted raw P3-minus-P2 contrast | 0.0952 | 0.0956 | deposited library, matched within time |
| Initial mouse tissue screen: primary exact within-time probability | 0.1111 | 0.1111 | 6 deposited libraries; 9 allocations |
| Initial mouse tissue screen: series with positive mean raw contrast | 2 / 2 | 2 / 2 | series; second is accession confounded |
| Initial mouse tissue screen: positive matched contrasts | 5 / 5 | 5 / 5 | 5 times nested within 2 dataset series |
| Depth-matched tissue audit: primary time-adjusted raw P3-minus-P2 contrast | 0.0494 | 0.0519 | AER-free source; 500-UMI pre-inference target |
| Depth-matched tissue audit: positive matched contrasts | 2 / 5 | 3 / 5 | scope boundary across 2 nested series |
| State-balanced macrophage mean of five raw contrasts | 0.01765 | 0.02217 | equal-state deposited-library summary |
| State-balanced macrophage positive matched contrasts | 5 / 5 | 5 / 5 | nested times; sign $P = 1/32$<br>corroborating |
| State-balanced macrophage primary exact probability | 0.1111 | 0.1111 | 6 deposited libraries; 9 blocked allocations |
| Complete-lineage screen: immune mean and positive contrasts | 0.03443 (5 / 5) | 0.03842 (5 / 5) | outcome-informed selection among 3 complete lineages |
| Complete-lineage screen: endothelial / mural positive contrasts | 2 / 5; 2 / 5 | 2 / 5; 2 / 5 | same depth-matched screen |
| Uniform full-space null: selection-adjusted empirical probability | 0.00010 | 0.00010 | 10,000 directions; lineage selection repeated per draw |
| Uniform full-space null: fixed macrophage empirical probability | 0.00020 | 0.00010 | 10,000 directions; mean of five raw contrasts |
| Source-covariance null: selection-adjusted empirical probability | 0.0820 | 0.0382 | 10,000 source-supported directions; lineage selection repeated |
| Source-covariance null: fixed macrophage empirical probability | 0.0970 | 0.0478 | 10,000 source-supported directions; fixed state-balanced summary |
| Source covariance retained by 5 principal components | 91.75% | 93.24% | 30 AER-free frog stratum-by-library centroids |
| Random directions with 5 / 5 positive contrasts | 12.52% | 12.75% | sign count alone is non-specific |
| Frog cell-cycle control: mean of five raw contrasts | 0.00668 (5 / 5) | 0.00301 (3 / 5) | AER-free cycling-minus-G1 ruler |
| State-balanced leave-one-frog-library-out rulers with positive five-contrast mean | 5 / 5 | 5 / 5 | 6 / 10 retain all five positive contrasts |
| Exact source-stage allocations with mean ≥ observed | 0.40 | 0.30 | all 10 allocations of three early / two late libraries |
| Joint tail-ROC permutation bridge probability | 0.0330 | 0.0410 | 1,000 within-library label shuffles |
| Reverse mouse ruler: frog limb stage 52 minus 58 | 0.03130; $P =$<br>0.00536 | 0.02808; $P =$<br>0.00357 | 8 deposited limb libraries; 560 stage allocations |
| Reverse mouse ruler: AER-free composition-standardized limb contrast | 0.02830; $P =$<br>0.10 | 0.03019; $P =$<br>0.10 | 5 deposited limb libraries; 10 stage allocations |
| Reverse mouse ruler: tail ROC-positive libraries | 11 / 14; $P =$<br>0.00330 | 13 / 14; $P =$<br>0.000183 | within-library ROC-minus-epidermis sign flips |
| Reverse mouse ruler: positive frog limb gaps after source omission | 6 / 6 | 6 / 6 | one complete mouse library omitted per ruler |
| Reverse mouse outcome labels: observed frog limb rank | 1 / 9 | 2 / 9 | all within-day outcome allocations |
| Skin wounded-at-PND2 minus wounded-at-PND21 raw contrast | 0.0382; $P =$<br>0.05 | 0.0411; $P =$<br>0.05 | 3 versus 3 libraries; all collected 7 days post wounding |
| Skin equal-lineage wounded-at-PND2 minus wounded-at-PND21 contrast | 0.0386; $P =$<br>0.05 | 0.0424; $P =$<br>0.05 | composition-reweighted deposited library |
| Skin equal-cell resamples positive | 1,000 / 1,000 | 1,000 / 1,000 | 300 cells per library per resample |
| Skin unwounded PND2-minus-PND21 raw contrast | 0.0461 | 0.0474 | contextual; 3 versus 3 deposited libraries |
| Skin descriptive wounded-minus-unwounded developmental contrast | -0.00785 | -0.00624 | collection ages differ across contexts; not an isolated injury effect |
| Canonical ten-seed source-target combinations positive | 100 / 100 | 100 / 100<br>separated | technical token-sampling audit |
| Ten-seed ensemble stage-52-minus-stage-58 raw contrast | 0.0611; $P = 0.0339$ | | deposited 5-dpa library |
| Homeolog-pair expression-effect correlation | $\rho = 0.483$; 7,702 pairs | | post hoc, equal-library expression audit |

*Scale note.* Unless a row is explicitly labelled as a probability, AUROC, correlation or standardized effect, decimal ruler results are raw, unitless differences between library-level summaries of internal cosine coordinates. They are not probabilities, percentages or calibrated amounts of regenerative competence. Standardized effects are raw differences divided by pooled between-library standard deviations and can exceed 1 in magnitude.

**Supplementary Table 3.** Targeted marker-set enrichment under an exact gene-label null.

| Previously defined marker set | Direction tested across frog and both mouse dataset series | Eligible / specified genes | Observed hits | Concordant background | Expected hits | Fold enrichment | Exact one-sided $P$ | Bonferroni $P$ |
| --- | --- | --- | --- | --- | --- | --- | --- | --- |
| Repair / lipid / phagocyte | positive in all three | 6 / 7 | 4 | 1,587 / 11,549 | 0.824 | 4.85 | 0.00423 | 0.00846 |

**Supplementary Table 4.**
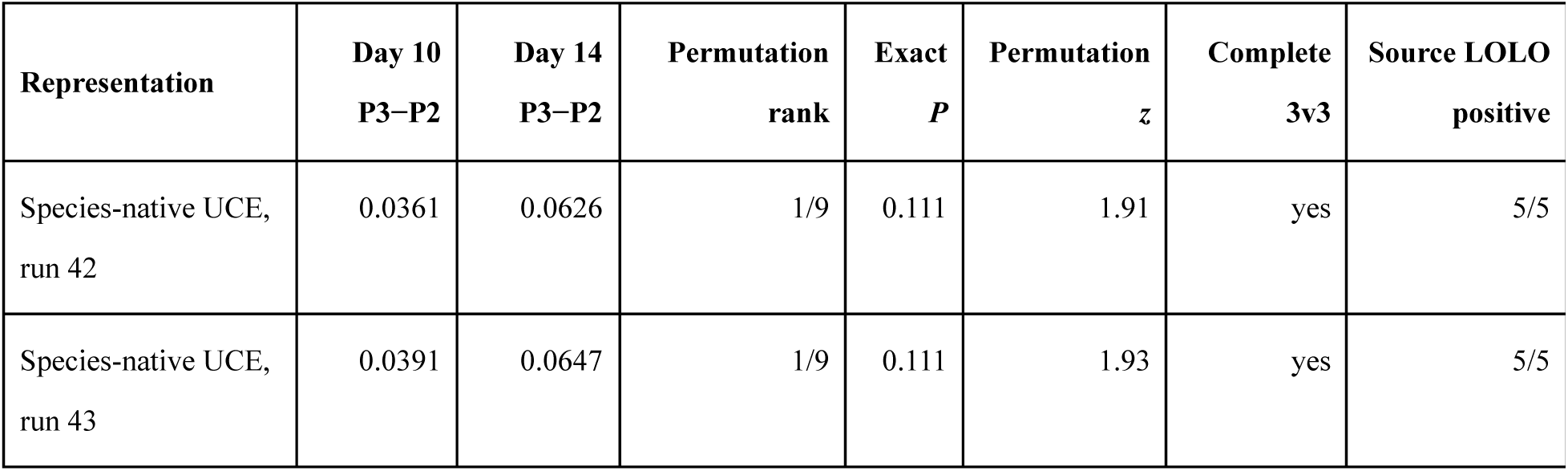
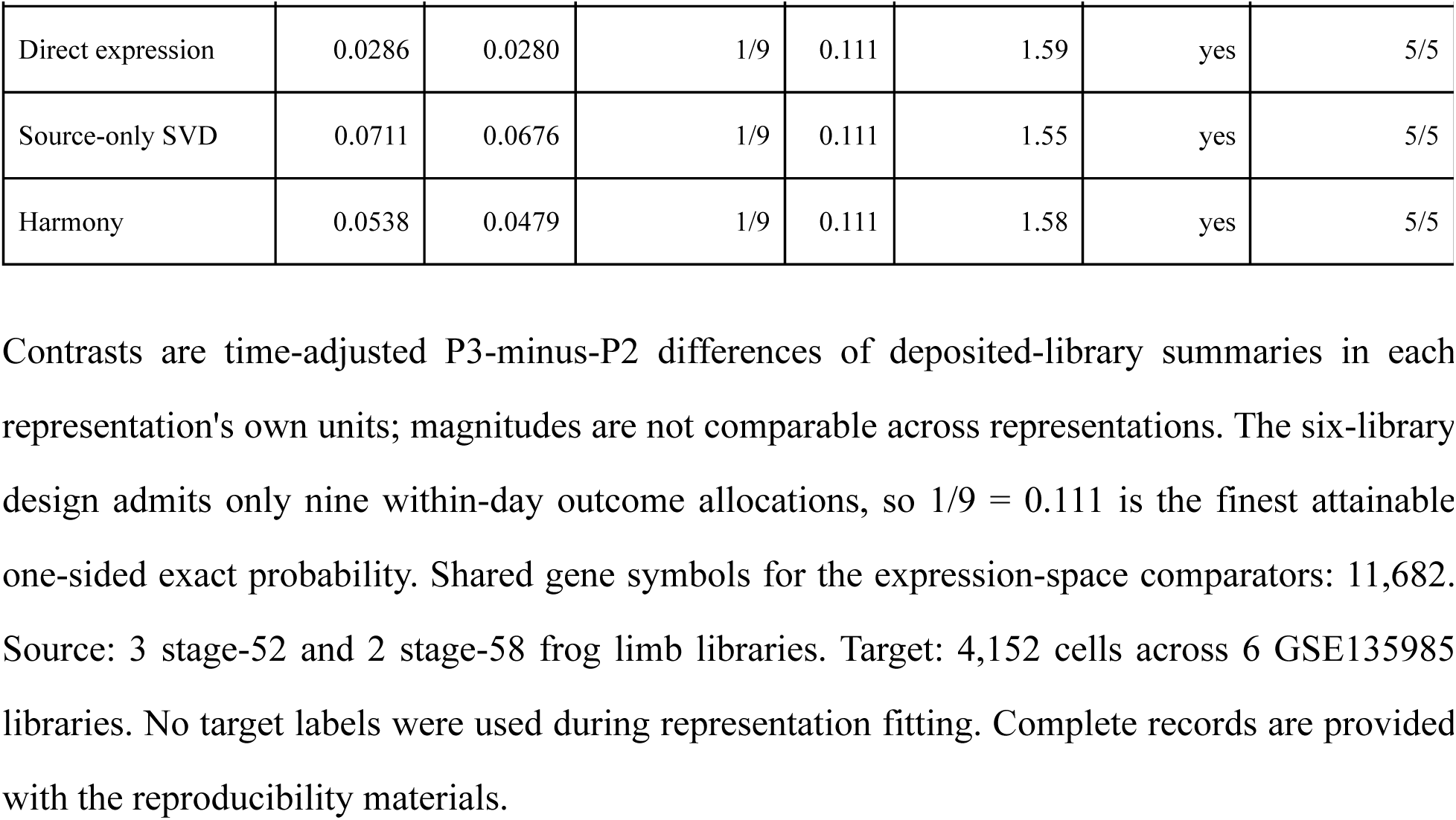
End-to-end representation comparison at the primary mouse digit endpoint.

| Representation | Day 10 P3–P2 | Day 14 P3–P2 | Permutation rank | Exact $P$ | Permutation $z$ | Complete 3v3 | Source LOLO positive |
| --- | --- | --- | --- | --- | --- | --- | --- |
| Species-native UCE, run 42 | 0.0361 | 0.0626 | 1/9 | 0.111 | 1.91 | yes | 5/5 |
| Species-native UCE, run 43 | 0.0391 | 0.0647 | 1/9 | 0.111 | 1.93 | yes | 5/5 |

| **Representation** | **Day 10**<br>**P3–P2** | **Day 14**<br>**P3–P2** | **Permutation**<br>**rank** | **Exact**<br>***P*** | **Permutation**<br>***z*** | **Complete**<br>**3v3** | **Source LOLO**<br>**positive** |
| --- | --- | --- | --- | --- | --- | --- | --- |
| Direct expression | 0.0286 | 0.0280 | 1/9 | 0.111 | 1.59 | yes | 5/5 |
| Source-only SVD | 0.0711 | 0.0676 | 1/9 | 0.111 | 1.55 | yes | 5/5 |
| Harmony | 0.0538 | 0.0479 | 1/9 | 0.111 | 1.58 | yes | 5/5 |

Contrasts are time-adjusted P3-minus-P2 differences of deposited-library summaries in each representation's own units; magnitudes are not comparable across representations. The six-library design admits only nine within-day outcome allocations, so $1/9 = 0.111$ is the finest attainable one-sided exact probability. Shared gene symbols for the expression-space comparators: 11,682. Source: 3 stage-52 and 2 stage-58 frog limb libraries. Target: 4,152 cells across 6 GSE135985 libraries. No target labels were used during representation fitting. Complete records are provided with the reproducibility materials.

